# Dissecting the cognitive phenotype of post-stroke fatigue using drift diffusion modeling of sustained attention

**DOI:** 10.1101/582502

**Authors:** Kristine M. Ulrichsen, Dag Alnæs, Knut K. Kolskår, Geneviève Richard, Anne-Marthe Sanders, Erlend S. Dørum, Hege Ihle-Hansen, Mads L. Pedersen, Sveinung Tornås, Jan E. Nordvik, Lars T. Westlye

## Abstract

Post-stroke fatigue (PSF) is a prevalent symptom among stroke patients. Its symptom burden is pervasive, persistent and associated with poor rehabilitation outcomes, though its mechanisms are poorly understood. Many patients with PSF experience cognitive difficulties, but studies aiming to identify cognitive correlates of PSF have been largely inconclusive. In contrast to conventional neuropsychological assessment, computational modeling of behavioral data allows for a dissection of specific cognitive processes associated with group or individual differences in fatigue. With the aim to zero in on the cognitive phenotype of PSF, we fitted a hierarchical drift diffusion model (hDDM) to response time data from Attention Network Test (ANT) obtained from 53 chronic stroke patients. The computational model accurately reconstructed the individual level response time distributions in the different ANT conditions, and hDDM regressions identified an interaction between trial number and fatigue symptoms on non-decision time, intuitively indicating that the cognitive phenotype of fatigue entails an increased vulnerability to sustained attentional effort. These novel results demonstrate the significance of considering the sustained nature of cognitive effort when defining the cognitive phenotype of post-stroke fatigue, and suggest that the use of computational approaches offers a further characterization of the specific processes underlying observed behavioral differences.

## Introduction

Post-stroke fatigue (PSF) is a common complaint among stroke survivors, with an estimated prevalence raging between 25 to 85% (Cumming, Packer, Kramer, & English, 2016). The symptom burden is often both pervasive and persistent (Duncan, Wu, & Mead, 2012; Schepers, Visser-Meily, Ketelaar, & Lindeman, 2006; van der Werf, van den Broek, Anten, & Bleijenberg, 2001) and associated with poorer outcome after rehabilitation, higher mortality (Michael, 2002; Naess, Lunde, Brogger, & Waje-Andreassen, 2012) and increased probability of institutionalization (Glader, Stegmayr, & Asplund, 2002). Post stroke fatigue has been defined as a highly prioritized future research topic by stroke survivors, family members and health care professionals (Pollock, St George, Fenton, & Firkins, 2012).

Although a universally accepted definition is lacking (Deluca, 2005), PSF is generally conceptualized as the feeling of debilitating tiredness and loss of energy (Stulemeijer, Fasotti, & Bleijenberg, 2005). Studies suggest that disruptions caused by the stroke lesion contribute to development of PSF (Naess et al., 2012; Wang, Wang, Wang, & Chen, 2014), though its etiological basis is likely multifactorial. Moreover, patients suffering from PSF often experience cognitive difficulties (Johansson & Rönnbäck, 2012; Pihlaja, Uimonen, Mustanoja, Tatlisumak, & Poutiainen, 2014), in particular when engaging in demanding activities over time, often referred to as mental or cognitive fatigue (Johansson & Ronnback, 2014). However, identifying robust and objective cognitive correlates of PSF has proven difficult, and although there is some evidence suggesting an association with attention, memory and processing speed (Lagogianni, Thomas, & Lincoln, 2018; Pihlaja et al., 2014), studies on the relationship between self-reported fatigue and cognitive function have been largely inconclusive (Holtzer, Shuman, Mahoney, Lipton, & Verghese, 2010; Ponchel, Bombois, Bordet, & Hénon, 2015). This may partly be due to the use of multifactorial neuropsychological tests, with varying or low cognitive specificity and which do not account for the temporal aspects during the course of a test session (Holtzer et al., 2010). With the assumption that a critical characteristic of cognitive fatigue is the failure to maintain or sustain cognitive effort over time, monitoring performance over time should increase sensitivity to cognitive manifestations of fatigue (Holtzer et al., 2010), and would also be closer in line with the conceptual definition of cognitive fatigue as “decreased performance during acute but sustained mental effort” (Deluca, 2005).

The attention network test (ANT) (Fan, McCandliss, Sommer, Raz, & Posner, 2002), combines a flanker test (Eriksen & Eriksen, 1974), and a cued reaction time task (Posner, 1980) in a computerized behavioral paradigm requiring sustained attention over time. The full version lasts for about 20 minutes, where accuracy and response times (RT) are tracked over time in 288 trials with varying cognitive demands. The ANT allows for estimation of individual level attention network scores such as the alerting, orienting and executive components, defined as relative differences in average RTs between different flanker and cue conditions (Fan et al., 2002). Although representing a widely applied and valuable contribution to theories on attentional function, analytical approaches based on mean RTs have suffered reliability problems, and are vulnerable to tradeoffs between speed and accuracy which are not accounted for in the model (Miller & Ulrich, 2013).

In contrast, computational approaches such as the drift diffusion model (DDM;(Ratcliff, 1978)) simultaneously models the full distribution of RTs and accuracies to estimate parameters reflecting specific theoretical cognitive constituents. DDMs are frequently applied to simple and speeded decision-making tasks (Ratcliff & McKoon, 2008; Ratcliff, Smith, Brown, & McKoon, 2016), offering both a theoretical framework to understand basic cognitive processes, and a psychometric tool to translate behavioral data into sub-components of cognitive processing (Ratcliff & McKoon, 2008). DDMs conceptualize decision-making as a noisy process where information is accumulated over time, continuing until a decision threshold is reached and a response is initiated (Ratcliff & McKoon, 2008). Four parameters are postulated in the original model (Ratcliff, 1978): *drift rate* (v), describing the rate or the speed of information accumulation, reflecting processing efficiency, *non-decision time* (t) representing time needed for stimulus encoding and response execution, *decision boundary separation* (a) indicating how much evidence is needed before a decision is made, and the *starting point* (z), reflecting any bias towards one of the two responses (Ratcliff & McKoon, 2008).

Applying computational models such as the DDM in clinical research may allow for a dissection of specific cognitive processes underlying observed group and individual differences in RT patterns. For example, assessing young and older subjects with a signal detection task, Ratcliff, Thapar, and McKoon (2001) found that the prolonged RTs often observed in older individuals were not explained by slower drift rates but rather longer non-decision times and higher decision thresholds, which provided a relevant adjustment to the long held notion of a general slowing in cognitive aging (Brinley, 1965; Salthouse, 1985).

Such computational approaches may thus offer a targeted assessment of cognitive function post stroke with higher sensitivity to self-reported fatigue than conventional neuropsychological assessment. To this end, we fitted a DDM to ANT behavioral data obtained from 53 chronic stroke patients and tested for associations between the model parameters (drift rate (v), non-decision time (t), and boundary separation (a)), and self-reported symptoms of fatigue as measured using the fatigue severity scale (FSS; (Krupp, LaRocca, Muir-Nash, & Steinberg, 1989)). To account for the temporal aspects of task performance, we specifically tested for interactions between FSS, trial number and performance. Based on current theories and the literature summarized above, we hypothesized that 1) individual differences in PSF burden would be associated with specific cognitive mechanisms as revealed by associations between FSS scores and DDM model parameters, and, 2) that any associations between PSF and model parameters will interact with trial number, with increasing associations between fatigue and model parameters with more sustained performance.

## Materials and methods

### Sample

Stroke patients who had been previously admitted with acute stroke to the Stroke Unit, Oslo University Hospital or the Geriatric department, Diakonhjemmet Hospital between 2013 and 2016, were invited by letter. Patients had to be in a chronic phase, defined as minimum 6 months post-stroke, with no other severe neurological, psychiatric or neurodevelopmental conditions. Among the approximately 900 invitation letters, 250 patients responded to decline or obtain more information. 77 were interested and eligible for inclusion and provided informed consent. 19 of the 77 patients withdrew during the course of the study, and four additional patients were excluded because of medical conditions. One additional patient was excluded due to behavioral criteria for the ANT (see below), resulting in a final sample of n=53 stroke patients.

Table 1 summarizes relevant demographic and clinical information of the patient group and Figure 1 shows the age distribution. This work was part of an intervention study on cognitive rehabilitation after stroke with a double baseline, randomized controlled design (see Kolskaar et al., 2019 for more details, including a description of overall study design). All data for the current study were collected from the baseline assessments prior to the intervention, starting 6 – 45 months after the acute stroke.

Mini-Mental State Examination (MMSE) scores <24 may indicate cognitive impairment and warrant further examination (Strobel & Engedal, 2008). One patient scored below 24, but further neuropsychological assessments done by a clinical psychologist indicated that cognitive function was sufficient for participation and that the inclusion criteria were not violated. The study was approved by the Regional Committee for Medical and Health Research Ethics, South-East Norway. All participants provided their written informed consent prior to inclusion.

**Table 1.**

| *Sample characteristics* |  |  |  |  |
| --- | --- | --- | --- | --- |
|  | Statistics | SD | Min | Max |
| *Current demographic and clinical information* |  |  |  |  |
| Age (mean) | 69.00 | 7.43 | 47 | 81 |
| Males/females (count) | 38/15 | - | - | - |
| Education in years (mean) | 14.56 | 3.65 | 9 | 30 |
| FSS (mean) | 3.53 | 1.46 | 1.11 | 6.77 |
| MMSE (mean) | 28.22 | 1.68 | 22 | 30 |
| *Stroke related information* |  |  |  |  |
| NIHSS at hospital discharge (mean) | 1.14 | 1.23 | 0 | 6 |
| TOAST classification for ischemic stroke* | Large artery atherosclerosis (19)<br>Small vessel occlusion (18)<br>Cardioembolism (6)<br>Other (10) |  |  |  |
| Months since stroke (mean) | 25.59 | 9.40 | 6.00 | 45.00 |

\*All but one patient suffered ischemic stroke

**Fig. 1.**
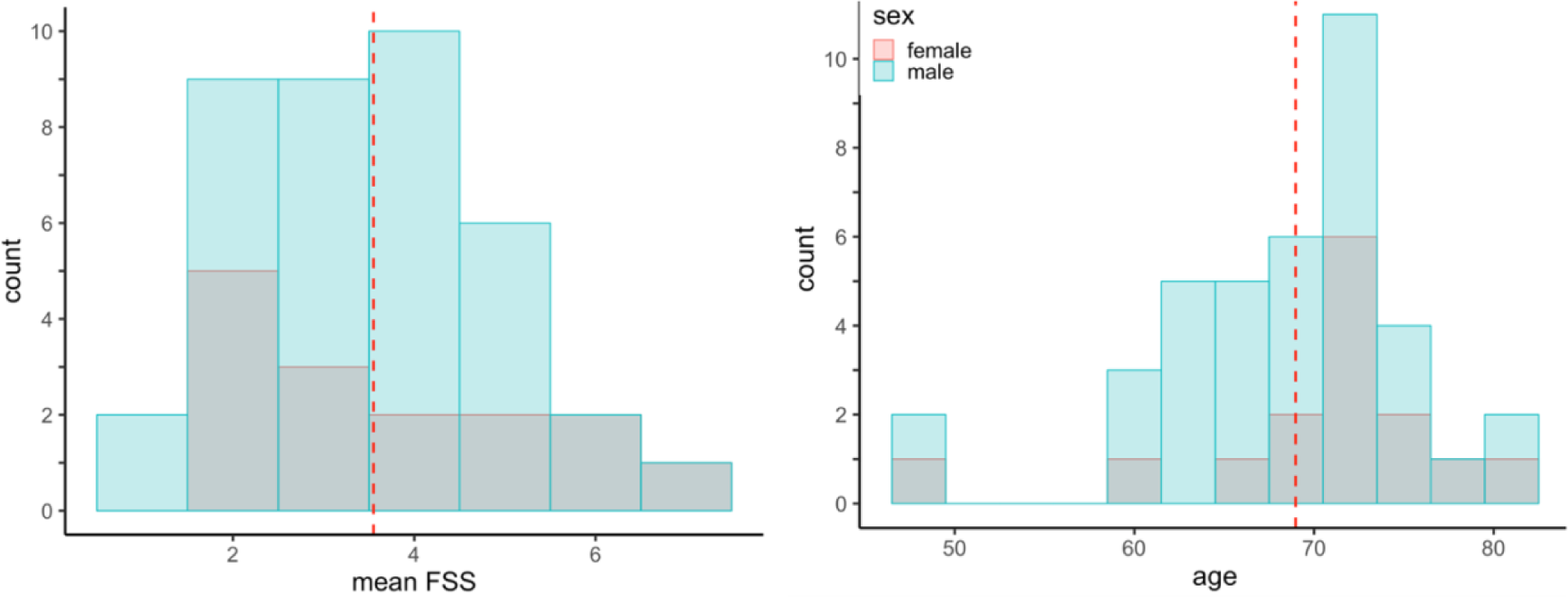
Distribution of sum FSS scores by gender and distribution of age by gender. Red line denotes the mean.

### FSS

Fatigue was measured by the FSS (Krupp et al., 1989), which is a one-dimensional, 9-item self-report scale, and one of the most frequently used measures to assess fatigue after stroke and other neurological conditions (Cumming, 2016; Lerdal et al., 2009; Whitehead, 2009). A review of 22 fatigue measures concluded that FSS was among the three scales that demonstrated good psychometric properties, as well as sensitivity to change in fatigue over time (Whitehead, 2009). Figure 1B shows the distribution of mean FSS scores. Average FSS score was 3.53 (SD=1.46), and 35% of the patients reported mean FSS>4, which is a commonly adapted threshold for clinical fatigue (Krupp et al., 1989; Schepers et al., 2006).

### Attention Network Test

A conventional version of the ANT was applied, as previously described (Fan et al., 2002). Figure 2 depicts the details of the task. Briefly, participants were instructed to keep their gaze towards a fixation cross (presented with a duration of 400, 800, 1200 or 1600 ms).

**Fig. 2.**
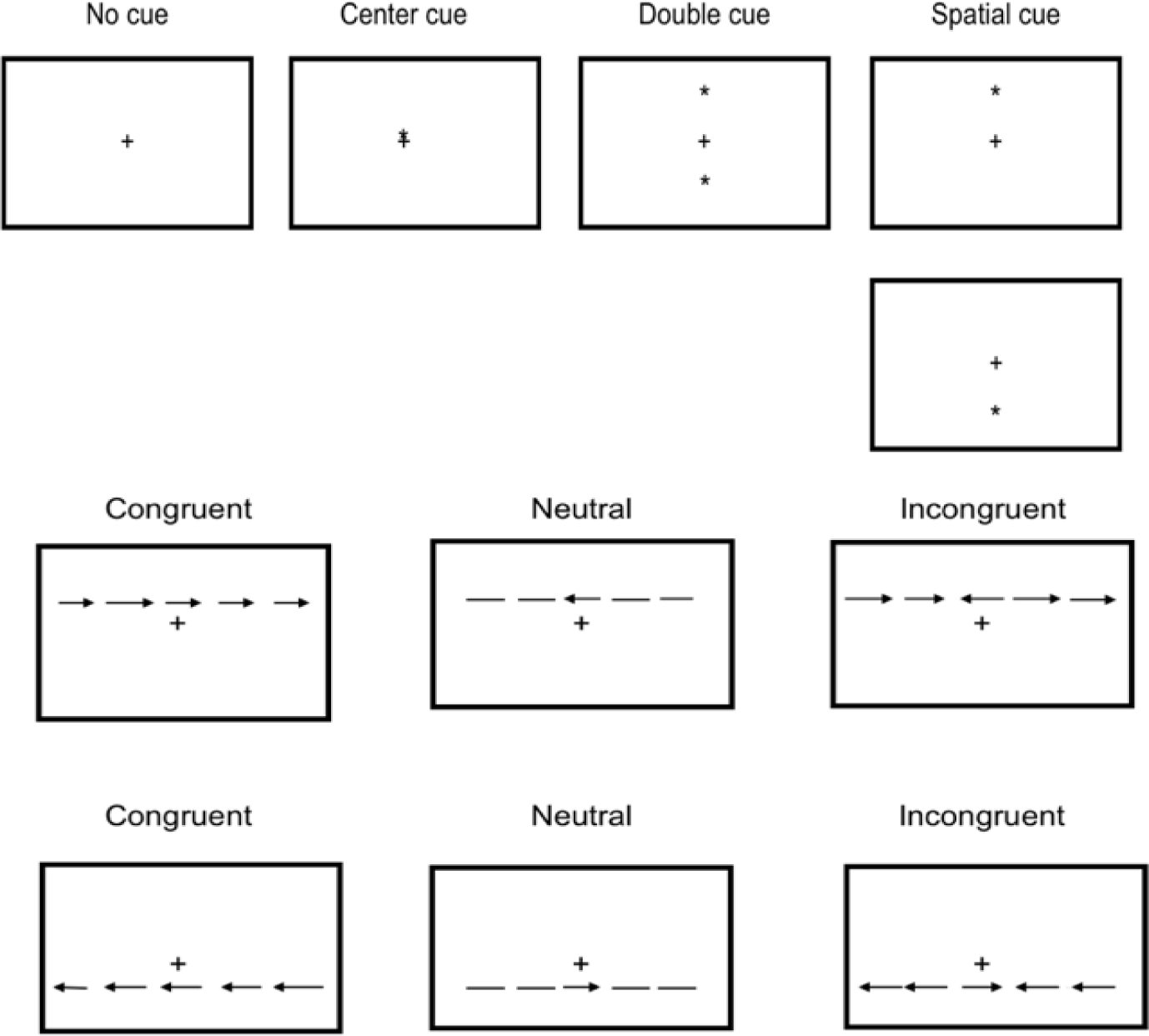
A schematic representation of the ANT cue and flanker conditions.

Immediately following the fixation cross, and preceding the target stimulus, one out of four cue conditions would appear for 100 milliseconds; a center cue (temporal cue), a double cue (temporal cue), a spatial cue (temporal and spatial cue) or no cue. Then, the task stimulus of five arrows were presented for 1700 milliseconds, and the participant had to decide whether the middle arrow (target arrow) were pointed left or right. Responses were executed by pressing the left or the right mouse button. The experimenter emphasized that responses should be made *as quickly and as accurately* as possible.

The middle target arrow was flanked by either congruent, incongruent or neutral arrows with regards to the direction of the target arrow. The incongruent, congruent and neutral flankers represent three different stimulus conditions, where the incongruent condition introduces a cognitive conflict (and thereby an executive component), typically resulting in longer reaction times and higher error rates (Westlye, Grydeland, Walhovd, & Fjell, 2010).

The task consisted of a practice block of 24 trials followed by 288 trials distributed equally over three rounds (96 trials per round). Each round lasted approximately 6 minutes. Participants were encouraged to take a short break between each round (maximum 2 minutes break). E-prime software (Psychology Software Tools, Pittsburgh, PA) were used for setting up the experiment and collecting the responses.

### Statistical analyses

#### Outlier exclusion and data cleaning

Trials with RT<200 ms, thought to reflect fast guesses, were removed from the analysis. 2% of the responses were removed due to this criterion. Participants having more than 50% incorrect responses within any of the flanker conditions were discarded. One participant was removed due to this criterion.

#### Hierarchical drift diffusion modeling

Cleaned RT and accuracy data were submitted to hierarchical drift diffusion modeling by use of the python toolbox HDDM (Wiecki, Sofer, & Frank, 2013). HDDM uses hierarchical Bayesian parameter estimation, which provides enhanced statistical power and allow for estimation of both individual and group parameters simultaneously (Wiecki et al., 2013). In addition to the cleaning describe above, an outlier mixture model included in the HDDM was applied, which assumes that a fixed proportion (5%) of trials are outliers that come from a uniform distribution not generated by a diffusion process (Wiecki et al., 2013). A mixed model allowing for some outliers has been shown to provide a better fit in likelihood models than models not allowing for any outliers at all (Wiecki et al., 2013).

#### Model selection / defining parameters

When parametrizing the DDM, we tested different cognitively plausible models to identify the model that best explained data, guided by the theoretical assumption that *drift rate (v)* should be allowed to vary as a function of stimulus difficulty condition (Ratcliff, Smith, & McKoon, 2015). Further, *decision threshold (a)* was assumed to be constant across stimulus conditions, following the logic that if *a* varies with stimulus conditions, the participant would have to first identify the condition, before adjusting threshold, and then start accumulating information from the stimulus, a sequence of events that does not seem plausible (Thapar, Ratcliff, & McKoon, 2003). *Non-decision time (t,* stimulus encoding and motor responses), was not expected to be affected by flanker condition, given that the visual stimuli were highly similar across flanker conditions and motor responses were simple and identical across conditions (simple button press).

Building on the abovementioned assumptions, we estimated different models and tested which combination of parameter fixations gave the best model fit, without providing any information about FSS scores (see Supplementary Table 1 for an overview of models tested). Finally, FSS scores were added to the best fitting regression model to estimate the effect of self-reported fatigue on model parameters and its interaction with time.

Model fit was assessed by comparing the deviance information criteria (relative DIC-values) between models. In Bayesian analyses, the DIC provides an estimation of fit of the model to the data, where lower DIC values indicate that the model has better support (François & Laval, 2011). As an additional test of model fit, we simulated data from the selected models and performed posterior predictive checks (PPC) to ensure that the model could reproduce central patterns in the observed data (Wiecki, 2016). 500 data sets were simulated by drawing 500 samples for each parameter from the estimated posterior distribution. The simulations thus capture the uncertainty in the estimated model, and allow for comparisons with the observed data.

#### Estimating the posterior distributions and assessing convergence (model diagnostics)

We used a Bayesian framework and Markov Chain Monte-Carlo sampling (MCMC) to estimate the posterior distributions (Kruschke, 2014). In the preliminary models, when testing and comparing parameter fixations, models were estimated on 1500 or 6000 samples. The final regression was run on 12000 samples. To improve convergence, the 4000 first samples were discarded, and thinning was set to 2 (keeping only every second sample).

A valid model should demonstrate convergence of the MCMC chains (Wiecki, 2016). Convergence was assessed by plotting and visually inspecting traces and autocorrelation plots for each estimated parameter. As a more formal test of convergence, the Gelman-Rubin statistics (R^; (Gelman & Rubin, 1992) was calculated. These values should be close to 1 and not exceed 1.1 if the chains have converged successfully, that is, if the samples of the different chains are similar (Wiecki et al., 2013).

#### Hypothesis testing within the hDDM

Effects of task and cue conditions, as well as the effects of time and fatigue status were determined by Bayesian hypothesis testing, by assessing the degree of overlap between posterior distributions. If less than 5 percent of the posterior distributions of two parameters overlap, the difference is said to be credible, or an effect is credibly different than null when at least 95 percent of the posterior distribution does not contain zero.

#### Associations between individual DDM parameters and FSS

To further explore the associations between task-related DDM parameters and subjective fatigue, individual model parameters (v, a and t) from DDM regression were extracted. Linear models in R, version 3.4.0 (RStudioTeam, 2016) were applied to test for associations between each of the model parameters and FSS scores while covarying for age and sex.

#### Associations between conventional ANT network scores and FSS

To compare the sensitivity of the DDM parameters with conventionally estimated ANT measures, we ran linear mixed models on each ANT network score and FSS scores. Based on a previous definition (Westlye et al., 2010), we computed the conventional ANT network scores orienting, alerting and executive control network based on median RTs and tested their relationship with FSS by linear regression models:

> Executive control = (RT incongruent – RT congruent) / RT congruent
>
> Alerting = (RT no cue – RT center cue) / RT center cue
>
> Orienting = (RT center cue – RT spatial cue) / RT spatial cue

#### Associations between median RT and FSS

Finally, in order to characterize the low-level dynamics in RT at the observed level, we used linear mixed models to test for main effects of FSS on RT, and for possible interactions between RT and trial number on FSS.

## Results

### hDDM regression models

The best fitting regression model allowed *drift rate (v)* to vary across flanker conditions, *non-decision time (t)* to vary across warning cue conditions and trial number while *boundary separation (a)* was kept constant (‘v ∼ flanker’, ‘t ∼ warningcue + trial’). The same regression was run with FSS added as a main effect and interaction term to t ∼ time (’v ∼ flanker, ‘t ∼ trial * FSS + warningcue’). The deviance information criteria (DIC) for the regression model including FSS was marginally better: −17648 with FSS versus −17645 without FSS. The variance parameter SV gave a slightly poorer fit (−17641), and was discarded. Posterior predictive checks indicated that the models managed to sufficiently reproduce observed patterns in the data (Supplementary Material). The regression model showed adequate convergence based on R-hat values, yielding two out of 660 Gelman-Rubin-statistics > 1.1 (1.102 and 1.16). Inspection of chains and autocorrelations confirmed adequate convergence for most parameters, but the standard deviations for t:FSS and t_time:FSS did not show optimal convergence (Supplementary Material).

### Effect of flanker conditions on drift rate

Figure 3 (left) shows the posterior probability plot for the drift rate estimated by flanker condition (intercept: congruent condition). The model provided strong evidence supporting that drift rate was lower in the incongruent condition compared to both congruent and neutral condition (P(v_Incongruent < v_Congruent) = 1.0, and P(v_Incongruent < v_Neutral) = 1.0), and drift rate was higher in neutral condition than in both congruent and incongruent condition (P(v_Neutral > v_Congruent) = 0.99, P(v_Neutral > v_Incongruent) = 1.0).

**Fig. 3.**
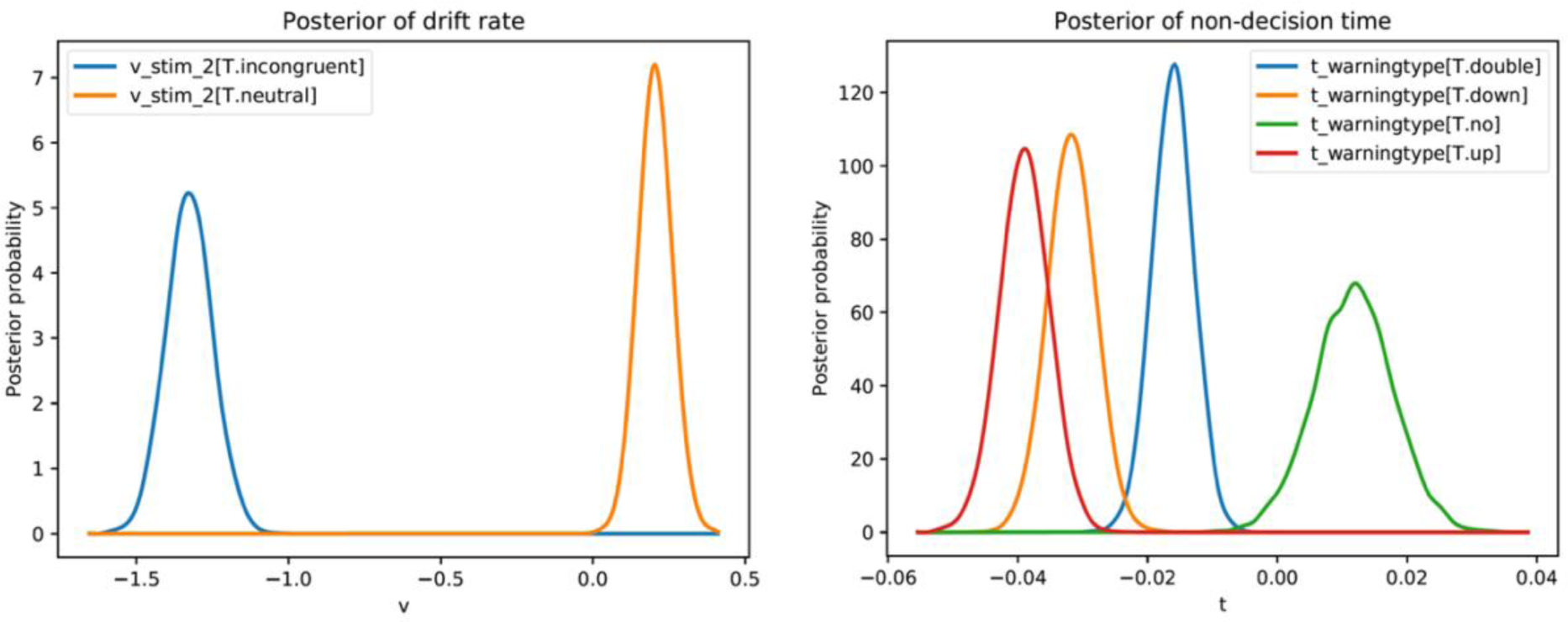
Posterior probability plot for drift rate (left) and non-decision time (right). Drift rate posteriors for incongruent and neutral flanker condition is plotted relative to the intercept (congruent flanker condition). Incongruent drift rate is lower than both congruent and neutral drift rate, and the distributions are not overlapping. In the right plot showing non-decision time, posteriors for different warning cue conditions is plotted relative to the intercept (center cue).

### Effect of warning cue on t, non-decision time

Figure 3 (right) shows the posterior probability plot for non-decision time (t) as a function of warning cue (intercept: center cue). Non-decision time was lowest for cue conditions “up” and “down”, and did not differ between these two cue conditions (P(t_down > t_up) = 0.92). “No cue” resulted in the highest non-decision time out of all cue conditions. “Center cue” (model intercept), gave the second-to highest non-decision time, lower than “no cue” (P(t_no > t_center) = 0.96), but higher than “double cue” (P(t_double < t_center) = 1.0).

### Effect of trial number and FSS on t, non-decision time

Figure 4 shows the posterior distributions for non-decision time. hDDM regression provided no support for a main effect of trial number on non-decision time (P(t_trial > 0) = 0.57) or of FSS (P(t_FSS > 0) = 0.39) on non-decision time. In contrast, the model provided evidence for an interaction effect between trial number and FSS on non-decision time (P(t_trial:FSS > 0) = 0.97), indicating that the association between FSS and non-decision time increases during the course of the experiment.

**Fig. 4.**
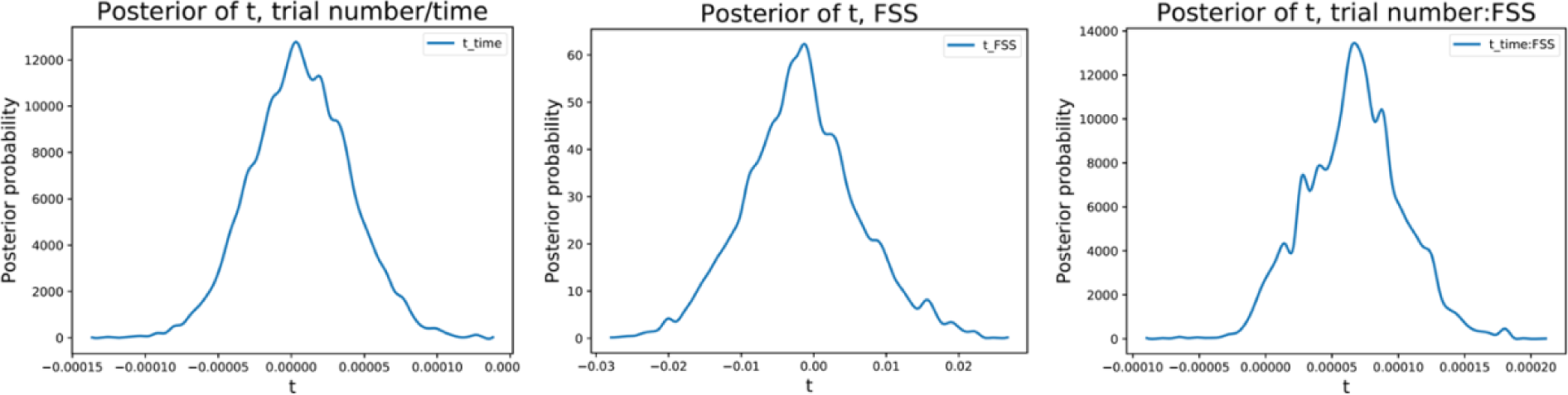
Posterior distributions of non-decision times (t) as a function of trial number (left) FSS score (middle), and the interaction between trial number and FSS (right). hDDM regressions provided no evidence in support of a main effect of trial number or FSS, but supported an interaction between FSS and trial number on t.

### ANT behavioral results

Supplementary Figure 1 shows RT distributions and means for correct responses in each flanker condition, across all subjects and cue conditions. Two-tailed t-tests revealed significant differences in RT between incongruent (M=776, SD=195) and congruent (M=666, SD=185, t = 28, p < 2.2e-16), between incongruent and neutral (M=654, SD=179, t = 32, p < 2.2e-16), and between neutral and congruent (t = −3.3, p = .0008).

There was no significant association between FSS and mean RT across (r=.09, p = 0.48) or within conditions (incongruent flanker: r = .05, p = 0.67, congruent flanker: r = .11, p = 0.47, neutral flanker: r = .12, p = 0.37). There was no association between number of errors and FSS across conditions or within the separate flanker conditions.

### Associations between ANT network scores and FSS and RT, trial number and FSS

Supplementary Figure 2 shows the distribution for each of the ANT network scores. One sample t-test revealed significant group level network score effects for Executive control network (mean=0.18, CI = 0.17 - 0.20, t = 21.57, p < 0.001), orienting network (mean = 0.06, CI = 0.05 – 0.08), t = 11.27, p < 0.001) and alerting network (mean = 0.04, CI = 0.03 – 0.06, t = 7.47, p < 0.001). Table 3 shows summary statistics from linear models testing for associations between ANT network scores and FSS. Whereas the analyses revealed a nominally significant association between FSS and the executive network score (t=-.23, p=.03), no associations remained significant after correction for multiple comparisons (corrected alpha = 0.05/6 number of tests = 0.008).

**Table 3.** Linear regression models by ANT network.

|  | Orienting |  | Alerting |  | Executive |  |
| --- | --- | --- | --- | --- | --- | --- |
|  | *t* | *p* | *T* | *p* | *t* | *p* |
| FSS | 0.57 | 0.567 | -1.03 | 0.305 | -2.23 | 0.030 |
| Age | 1.10 | 0.276 | -2.17 | 0.034 | -1.18 | 0.243 |
| Sex | 1.66 | 0.103 | -0.41 | 0.681 | -0.03 | 0.975 |

Table 4 shows the summary statistics from linear mixed-effects models testing for associations between RT and FSS, trial number, sex, and age, as well as the interaction between FSS and trial number, within each flanker condition. Figure 5 shows the estimated RT for each flanker condition for each group (low versus high fatigue, conveniently based on a median split for visualization purposes. See supplementary figure 5 for raw RT data). Briefly, after correction for multiple comparisons (corrected alpha = 0.008) the interaction between trial number and FSS was significant in the neutral and incongruent condition, as was the association between age and RT. There was no significant main effect of FSS on RT in any condition.

**Table 4.** Linear mixed-models by flanker stimulus

|  | Neutral |  | Incongruent |  | Congruent |  |
| --- | --- | --- | --- | --- | --- | --- |
|  | *t* | *p* | *T* | *p* | *t* | *p* |
| FSS | 0.65 | 0.516 | 0.11 | 0.911 | 0.84 | 0.403 |
| Trial | -1.66 | 0.096 | -1.62 | 0.105 | -0.11 | 0.907 |
| Trial:FSS | 4.11 | <0.001 * | 4.11 | <0.001* | 2.41 | 0.015 |
| Sex | -0.11 | 0.907 | -0.01 | 0.984 | 0.03 | 0.970 |
| Age | 3.66 | <0.001* | 3.16 | 0.002* | 2.41 | 0.001* |

\**p*-values remained significant after Bonferroni correction for multiple comparisons

**Fig. 5.**
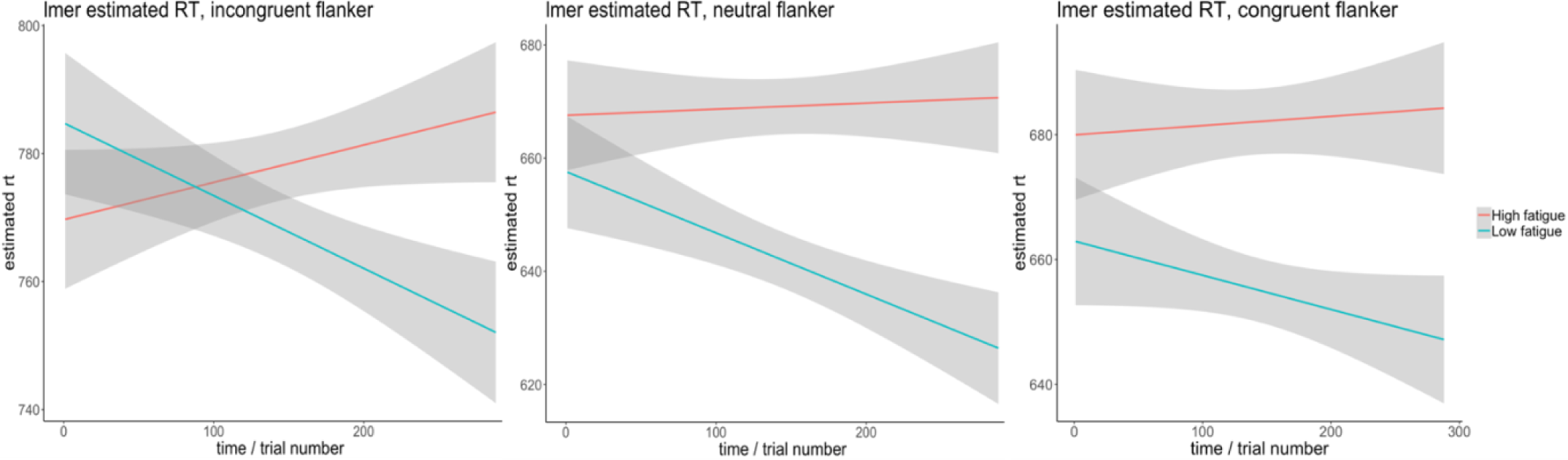
Estimated RT from linear mixed-effects models plotted by group (high/low fatigue, median split).

## Discussion

Post-stroke fatigue is a common and debilitating symptom in stroke patients, yet the brain and cognitive mechanisms are not well understood. Previous studies attempting to characterize the cognitive phenotype of PSF have reported inconsistent findings, possibly due to the methodological variability and the use of instruments lacking cognitive sensitivity and specificity. Computational modeling approaches applied on attentional behavioral tasks may provide a more targeted assessment of cognitive function post stroke with higher sensitivity to self-reported fatigue than conventional neuropsychological assessment. To this end, we fitted a hierarchical drift diffusion model to ANT response time data from 53 stroke patients. We hypothesized that individual differences in PSF burden would be associated with specific cognitive mechanisms as revealed by associations between FSS scores and DDM model parameters, and, that any associations between PSF and model parameters would interact with the duration of the effort, with increasing associations between fatigue and model parameters after sustained performance.

hDDM regression revealed no main effect of FSS on any of the model parameters, but provided evidence of an interaction between trial number and FSS on non-decision time, indicating increasing effects of FSS during the course of the experiment. This is in line with the second hypothesis, suggesting stronger associations between fatigue and DDM parameters with sustained performance, and demonstrates that hDDM is sensitive to fatigue in a cognitive context when explicitly modeling the interactions with time.

The relationship between sustained performance and fatigue was further supported by linear regressions on RT data, identifying significant interactions between fatigue and trial number on RT in the neutral and incongruent flanker conditions. In general, while patients with low levels of fatigue showed decreased RT over time, possibly reflecting learning effects, patients with high levels of fatigue showed slightly increased Rt during the course of the experiment. Analyses based on observed data alone do not allow for any inference regarding the specific cognitive processes that may underpin this effect of fatigue on RT. In contrast, the current computational dissection of the ANT data provided by hDDM indicated that the interaction between fatigue and trial number on RT was best explained by non-decision time, and not the speed of evidence accumulation (drift rate) or response style (boundary separation).

These results provide a novel account of the specific cognitive underpinnings of PSF. When the task context is appropriate, DDM parameters can be interpreted directly (Froehlich et al., 2016) and thus provide insight into the modular and temporal evolution of the decision process. Decision boundary separation (a) adjusts the tradeoff between speed and accuracy (Pedersen, Frank, & Biele, 2017). Large estimates of (a) is typically interpreted as indicative of a conservative decision style, associated with higher RTs but more accurate responses (Pedersen et al., 2017). Larger estimates of drift rate (v) are typically interpreted as more efficient information processing, and are expected to vary by “the quality of the information extracted from the stimulus” (Ratcliff & McKoon, 2008, p. 3), implying that experimental conditions varying in difficulty should produce different drift rates (Ratcliff & McKoon, 2008). In line with this, DDM identified a credible effect of flanker type on drift rate, with the more cognitively demanding incongruent condition resulting in the lowest drift rate, while neutral flankers yielded the highest drift rate.

Further, non-decision time (t) comprises both sensory encoding and motor response output (Ratcliff & Smith, 2010). The analyses revealed no main effect of trial number or FSS on non-decision time, but an interaction between trial number and FSS on non-decision time, indicating higher increases in non-decision time during the course of the experiment in patients with more fatigue symptoms. The lack of associations with drift rate and boundary separation indicates that fatigue is specifically associated with non-decision aspects of the response process, such as stimulus encoding or response execution rather than with the speed or efficiency of the evidence accumulation or with the decision threshold (i.e. how much information is required before making a decision). Previous studies have reported higher non-decision times in older compared to younger individuals (Ratcliff et al., 2001), and in this respect, patients reporting high fatigue are responding more like elderly individuals, but only after sustained exertion.

Research aiming to delineate the nervous system pathophysiology of PSF may further inform hypotheses about this apparent link between non-decision time and fatigue. Applying transcranial magnetic stimulation (TMS), a previous study (Kuppuswamy, Clark, Turner, Rothwell, & Ward, 2014) reported higher motor thresholds in stroke patients with high fatigue, and suggested that patients with PSF experience diminished excitability of motor pathways, regarding both corticospinal outputs and facilitatory inputs. In this respect, the current observation of an interaction between trial number and fatigue on non-decision time might reflect altered neuronal excitability. However, how such neurophysiological mechanisms would translate into the subjective perception of fatigue remains unclear. Here, the perception of effort might be central, in the sense that subjective fatigue may manifest when volitional motor cortex input does not longer produce the expected output due to reduced excitability (Kuppuswamy et al., 2014).

The coping hypothesis (Van Zomeren, 1984; Van Zomeren & Van den Burg, 1985) offers another explanatory framework for the observed interaction between sustained effort and FSS on non-decision time. Originally articulated in relation to traumatic brain injury patients, this view suggests that the chronic effort needed to compensate for cognitive deficiencies gives rise to secondary symptoms, hereunder fatigue. Hence, subtle cognitive deficits associated with stroke may be temporarily disguised by a compensating and temporary increase in cognitive effort. However, this compensation comes with the cost of increased feeling of fatigue, in particular during sustained effort. In line with this, the current interaction between trial number and fatigue may be understood as a result of increased cognitive effort, producing increased tiredness over time, resulting in a decline in performance. Future studies assessing brain activation patterns using functional brain imaging may help clarify the putative cost of compensation during cognitive effort in patients with fatigue.

Some limitations should be considered when interpreting the results of the current study. In line with most clinical studies, the study design does not allow for causal inference. Still, the findings may pave the way for future clinical or experimental studies examining possible causal mechanisms, and subsequent interventions targeting a specific component of the extended cognitive phenotype of PSF. Further, fatigue is a complex and multifactorial syndrome that often co-occur with depression (Høgestøl et al., 2018), which represents a challenge for all research aiming to target unique and specific characteristics of fatigue. Commonly applied depression questionnaires assess complaints that are also central to fatigue (i.e. feeling tired or having little energy, sleeping too much, trouble concentrating), and correlations between fatigue and depression measures are therefore expected. Hence, we did not adjust for depression scores in our analyses since this would imply a risk of removing much of the fatigue effects in our data together with the depression (Finke et al., 2015), and larger datasets are required for parsing the common and unique features among these intertwined conditions. Moreover, fatigue is more frequent than depression in stroke patients and many patients report fatigue without suffering from depression (Glader et al., 2002; van der Werf et al., 2001), and depression do not provide a simple account of the heterogeneity in PSF prevalence between studies (Cumming et al., 2016).

The distribution of NIHSS scores indicates that the current patients were sampled from a relatively healthy part of the full population of stroke patients. It is possible that a higher fatigue symptom burden on the group level could reveal associations that were not expressed in this relatively well functioning patient sample. Reported fatigue levels are, however, comparable with what has been reported in other studies (Wang et al., 2014), and higher than what is reported in healthy control samples (Valko, Bassetti, Bloch, Held, & Baumann, 2008). Further studies are needed to test the generalizability of the findings to different and more severely affected patient samples.

Although ANT appears to be a suitable paradigm to target cognitive aspects of PSF, as performance requires sustained attentional and executive resources (Holtzer et al., 2010), it should be noted that the error rate in the sample was low. This might have implications for the validity of the results from hDDM regression, because the model estimates parameters based on distributions of both RT and accuracy, and assumes different RT distributions for correct versus erroneous responses. One way this might have influenced the results, is that a relatively large proportion of the variation in the data may have been attributed to non-decision time, because this parameter is not assumed to be affected by accuracy (in contrast to v or a, which both vary with accuracy). The models should therefore be replicated in a larger sample with higher error rates. However, it is encouraging that the model performs reasonably well in predicting the few errors that are present in the data, indicating that the results are valid despite low error rates.

In conclusion, the current study represents a novel approach to dissect the cognitive phenotype of fatigue in stroke patients. The application of an advanced computational model on the temporal evolution of response times during the course of a demanding and sustained attentional task enabled the possibility to parse the observed response time patterns into specific cognitive processes. Specifically, the results indicate a relationship between the subjective experience of fatigue and response time distributions from a sustained attention task, and demonstrates the significance of considering the sustained nature of the task when targeting fatigue in a neuropsychological context, intuitively indicating that the cognitive phenotype of fatigue entails an increased vulnerability to sustained effort. In general, the use of computational approaches in the neuropsychological workup may offer a dissection of the specific cognitive processes underlying observed behavioral differences, with great clinical relevance.

## Acknowledgements and funding

This study was supported by the Norwegian ExtraFoundation for Health and Rehabilitation (2015/FO5146), the Research Council of Norway (249795, 262372), the South-Eastern Norway Regional Health Authority (2014097, 2015044, 2015073), and the Department of Psychology, University of Oslo.

