## Supplementary Material for "Dissecting the cognitive phenotype of post-stroke fatigue using drift diffusion modeling of sustained attention"

| Model | Samples | DIC |
|---|---|---|
| hddm.hDDMregression(a ~ warningcue) | 1500 | -15424 |
| hddm.hDDMregression(t ~ warningcue) | 1500 | **-15717** |
| hddm.hDDMregression(v ~ warningcue) | 1500 | -15169 |
| hddm.hDDMregression(a ~time) | 1500 | -14.356 |
| hddm.hDDMregression(t ~time) | 1500 | **-14.448** |
| hddm.hDDMregression(v ~time) | 1500 | -14.219 |
| hddm.HDDMRegressor(data, ['v ~ flanker, 't ~ time + warningcue'], group_only_regressors = False, p_outlier = 0.05) | 6000 | -17645 |
| **hddm.HDDMRegressor(data, ['v ~ flanker, 't ~ time * FSS + warningcue'], group_only_regressors = False, p_outlier = 0.05)** | 6000 | **-17648** |
| hddm.HDDMRegressor(data, ['v ~ flanker, 't ~ time * FSS + warningcue'], group_only_regressors = False, p_outlier = 0.05, include = 'sv' ) | 6000 | -17641 |

**Supplementary Table 1.** Model fits for various hDDM regression models. DIC: deviance information criterion, where lower values indicate a better model fit.

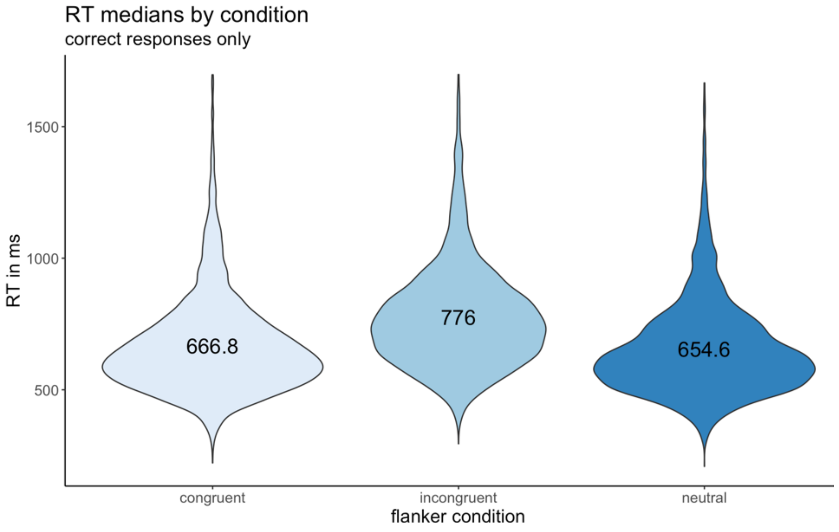


**Supplemental Fig. 1.** Reaction time distribution with medians for each flanker condition. Only correct trials included.

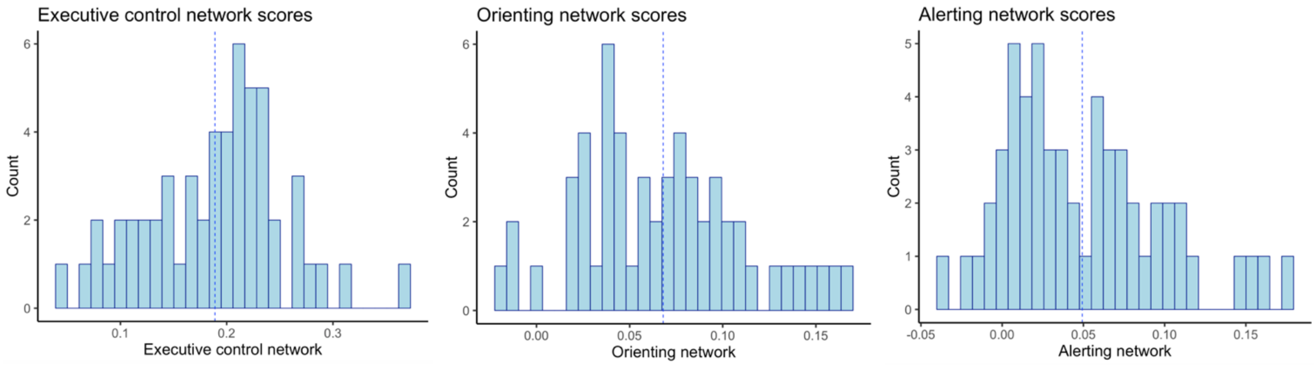


**Supplementary Fig. 2.** Histograms of ANT network scores. Left to right: Executive control network, orienting network and alerting network. Mean network score denoted by blue line.

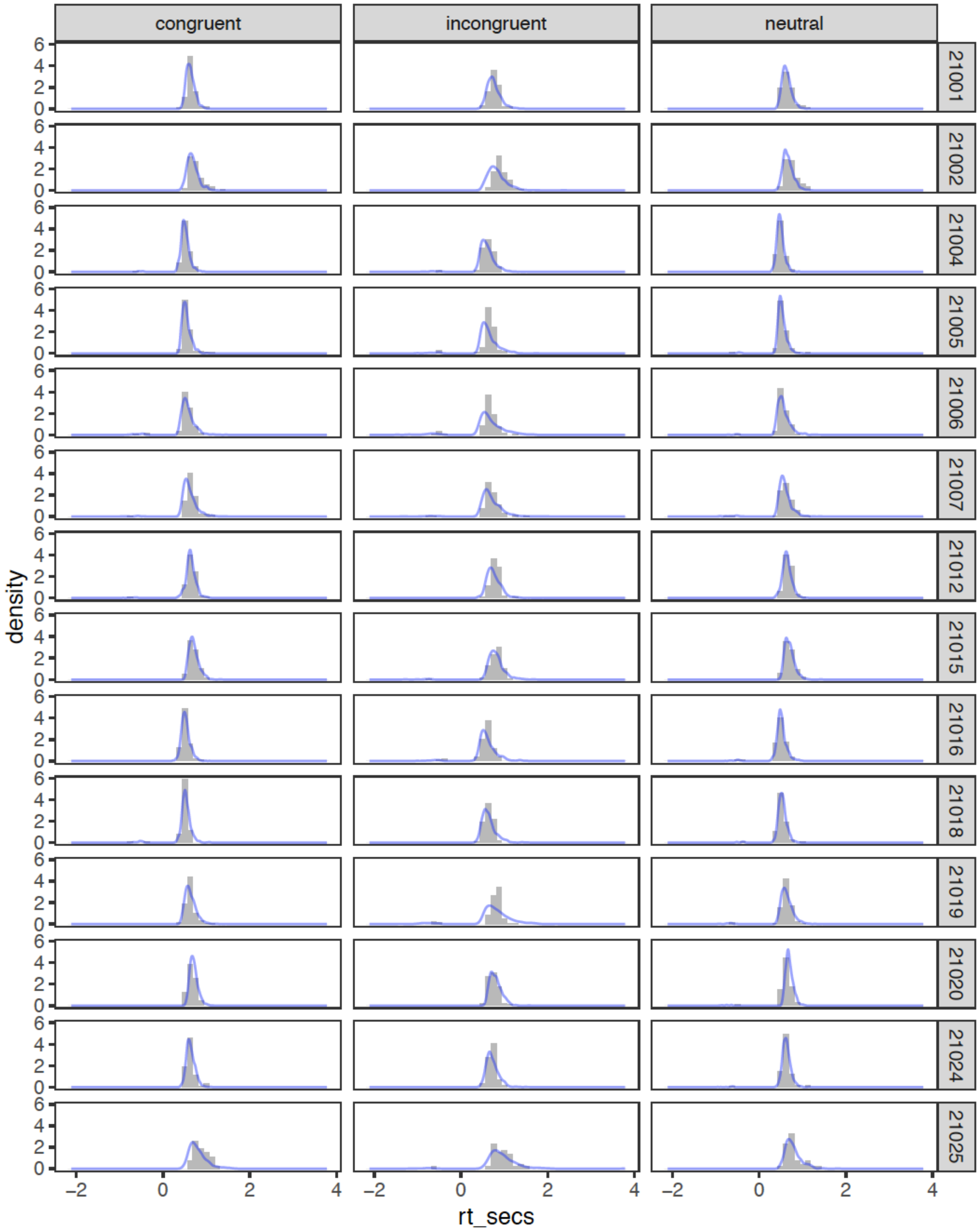


**Supplementary Fig. 3.** Posterior predictive plots. Response time histogram on the posterior predictive of the model (blue) for each patient. Correct trials are plotted as a positive distribution while errors are shown as negative.

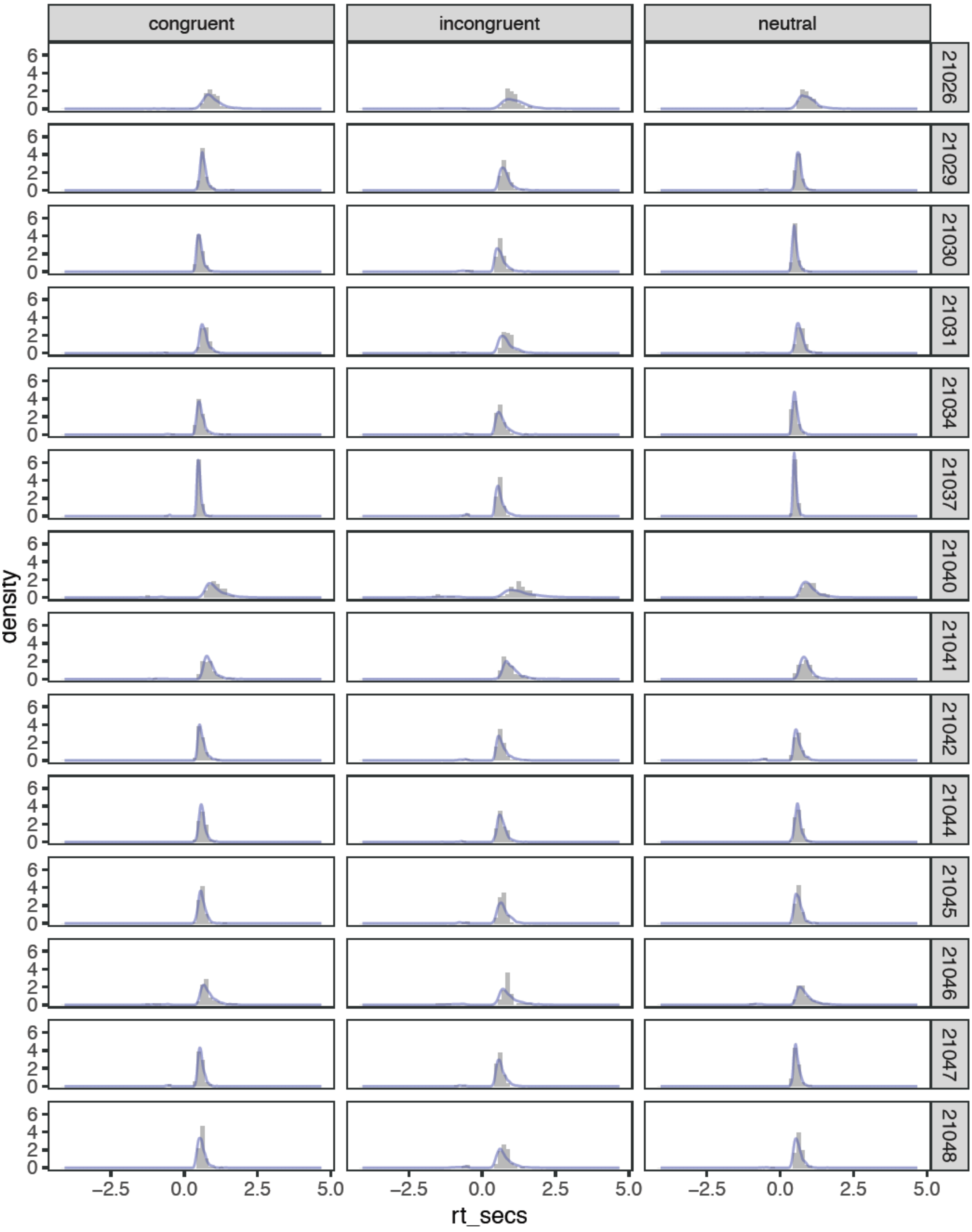


**Supplementary Fig. 3 (cont.).** Posterior predictive plots. Response time histogram on the posterior predictive of the model (blue) for each patient. Correct trials are plotted as a positive distribution while errors are shown as negative.

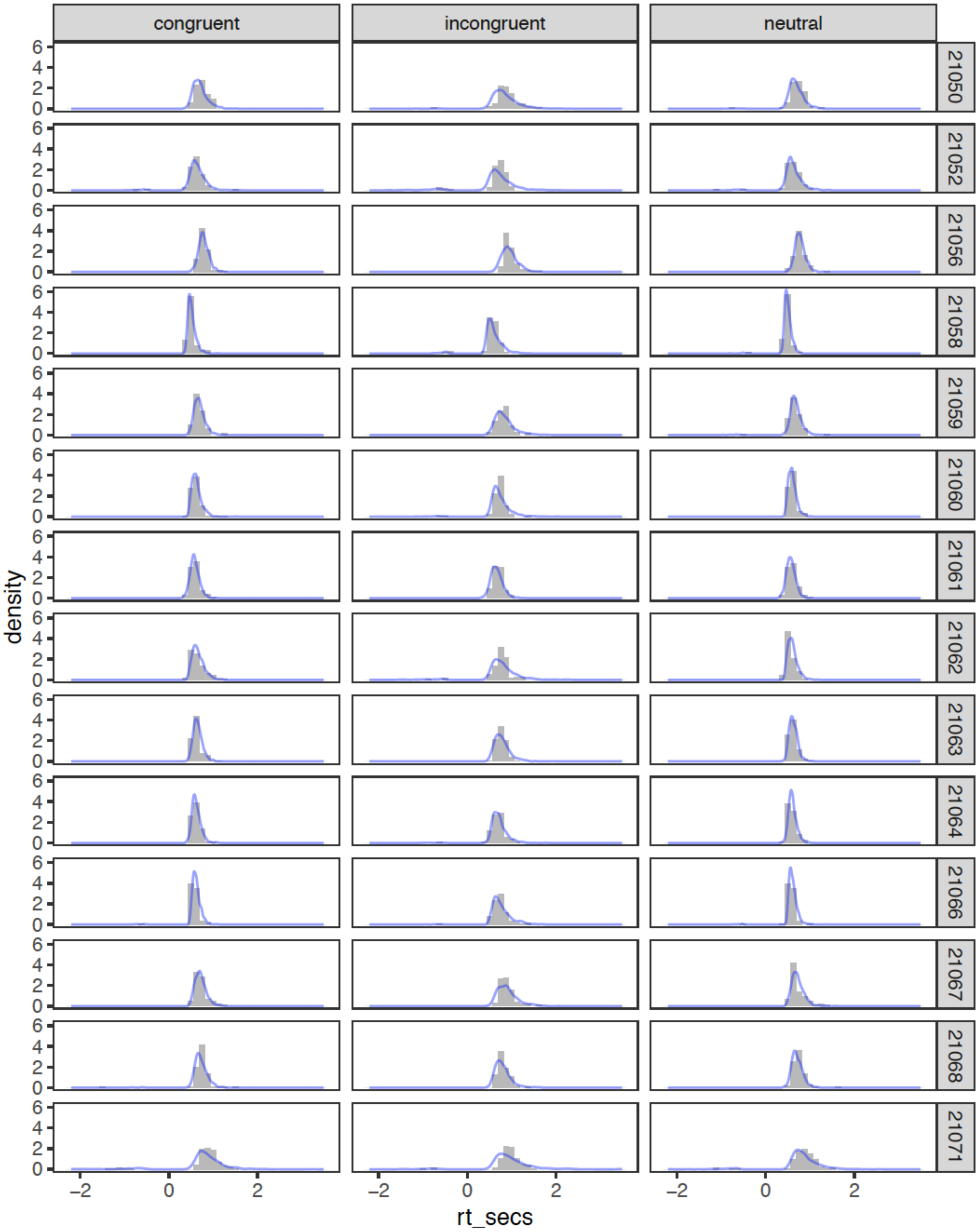


**Supplementary Fig. 3 (cont.).** Posterior predictive plots. Response time histogram on the posterior predictive of the model (blue) for each patient. Correct trials are plotted as a positive distribution while errors are shown as negative.

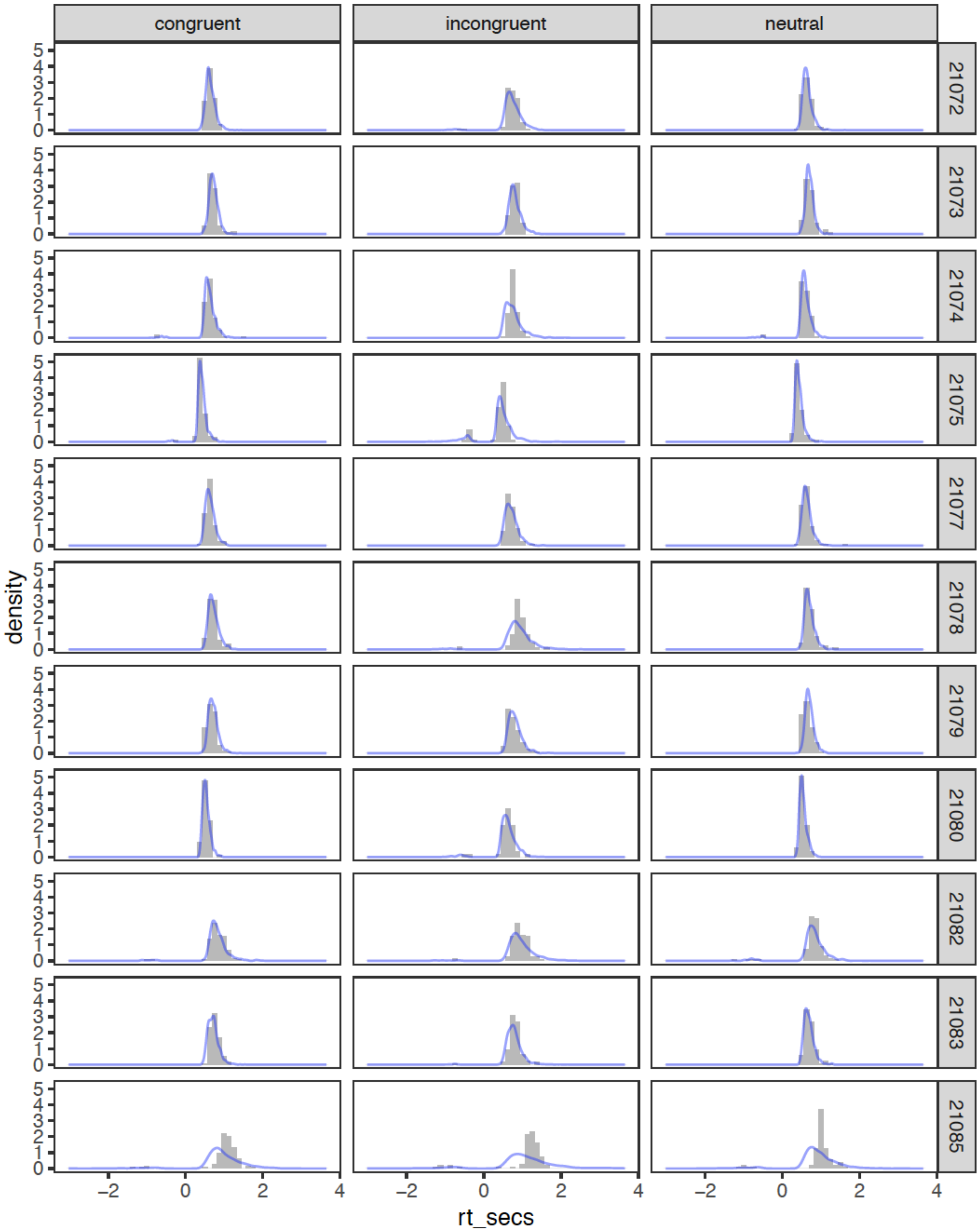


**Supplementary Fig. 3 (cont.).** Posterior predictive plots. Response time histogram on the posterior predictive of the model (blue) for each patient. Correct trials are plotted as a positive distribution while errors are shown as negative.

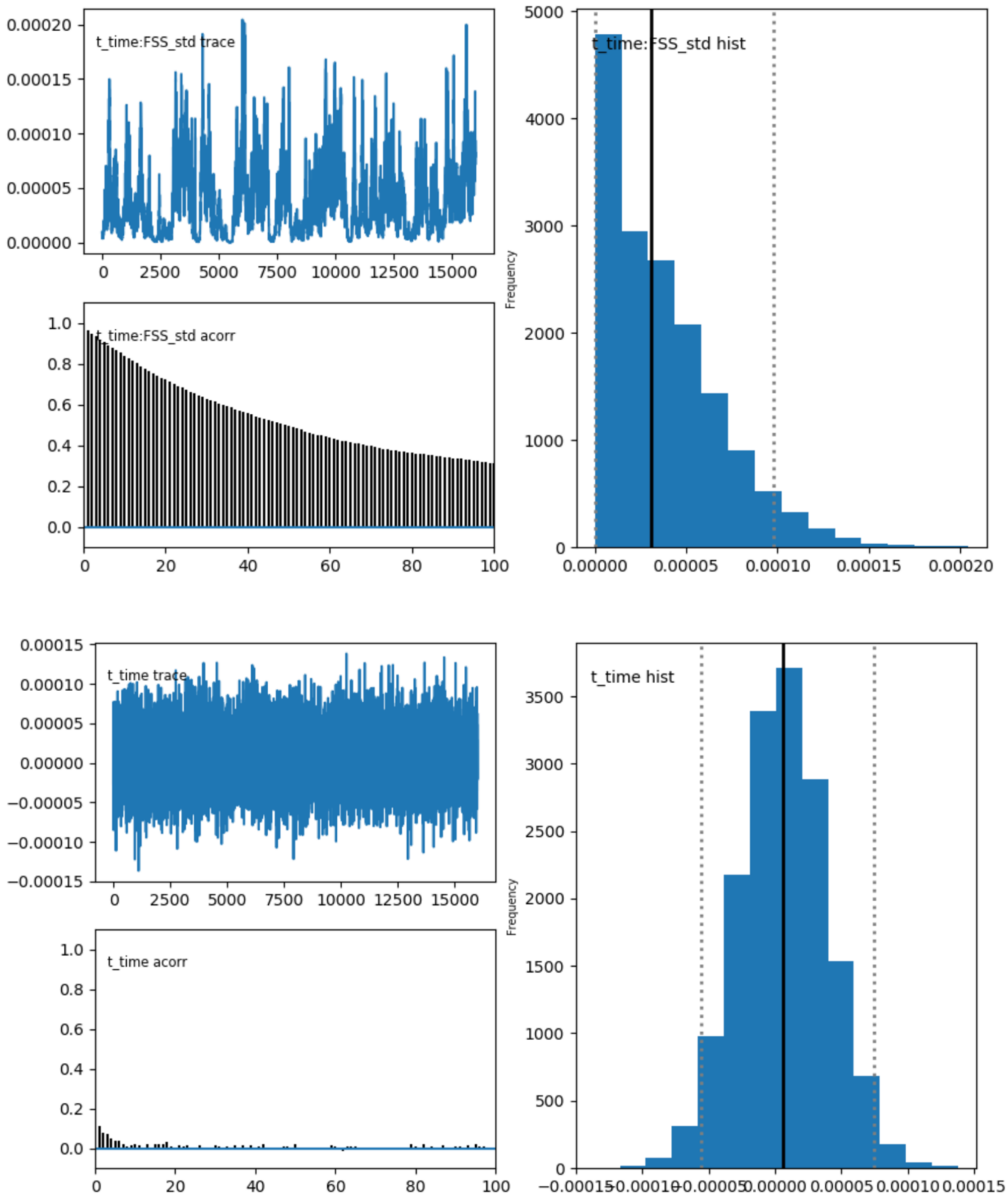


**Supplementary Fig. 4.** Traces, autocorrelations and histograms of all parameters included in the hDDM regression model.

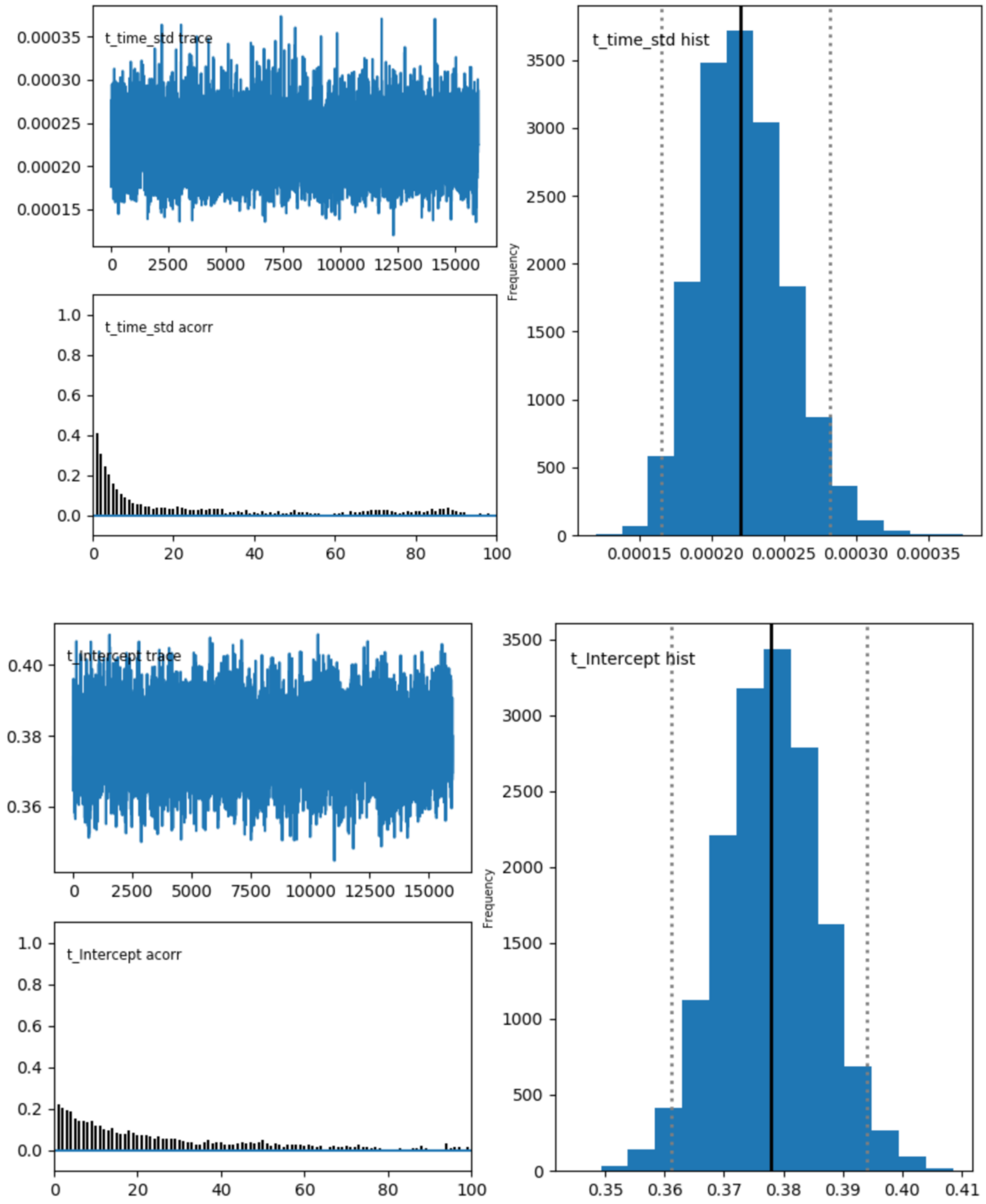


**Supplementary Fig. 4 (cont.).** Traces, autocorrelations and histograms of all parameters included in the hDDM regression model.

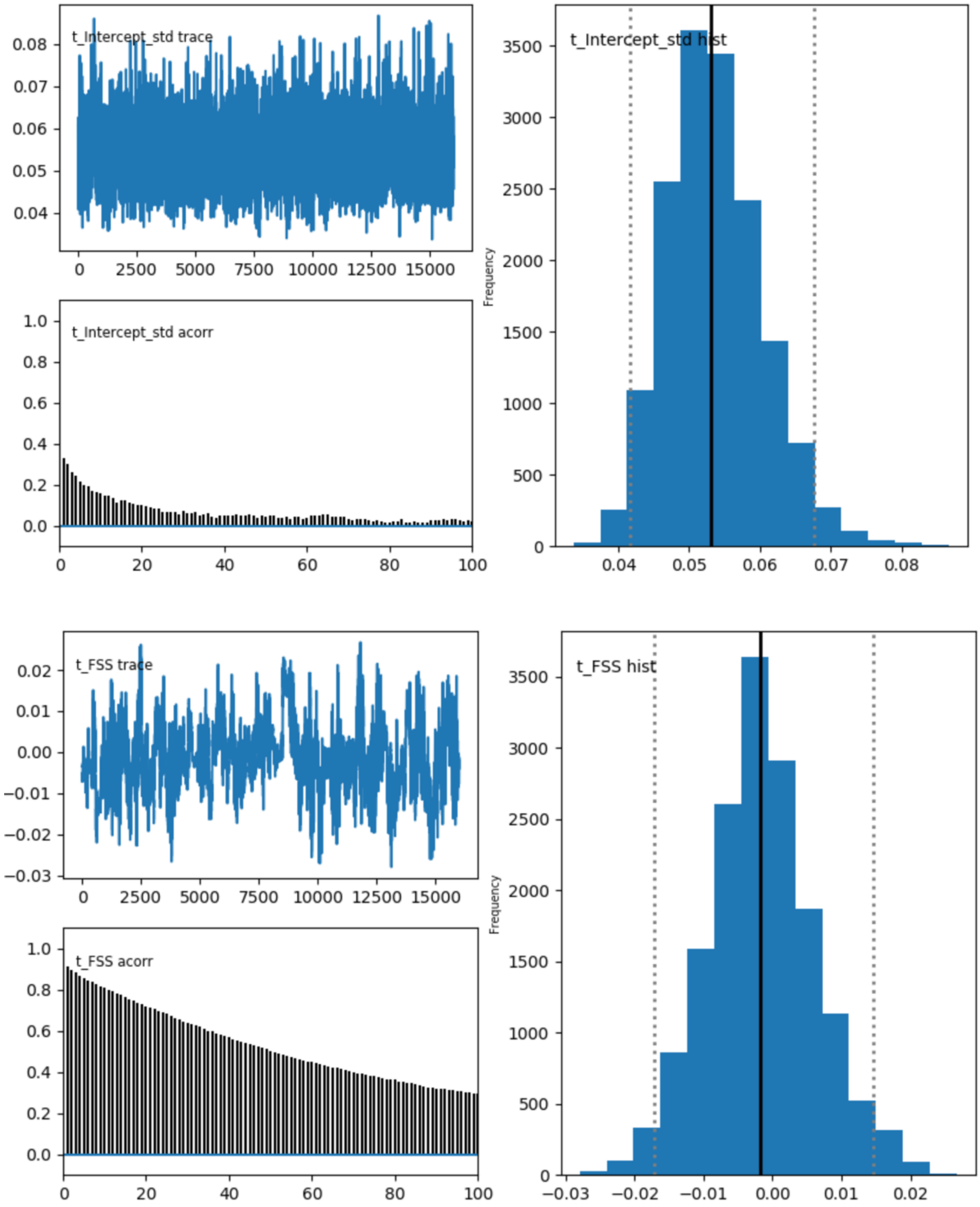


**Supplementary Fig. 4 (cont.).** Traces, autocorrelations and histograms of all parameters included in the hDDM regression model.

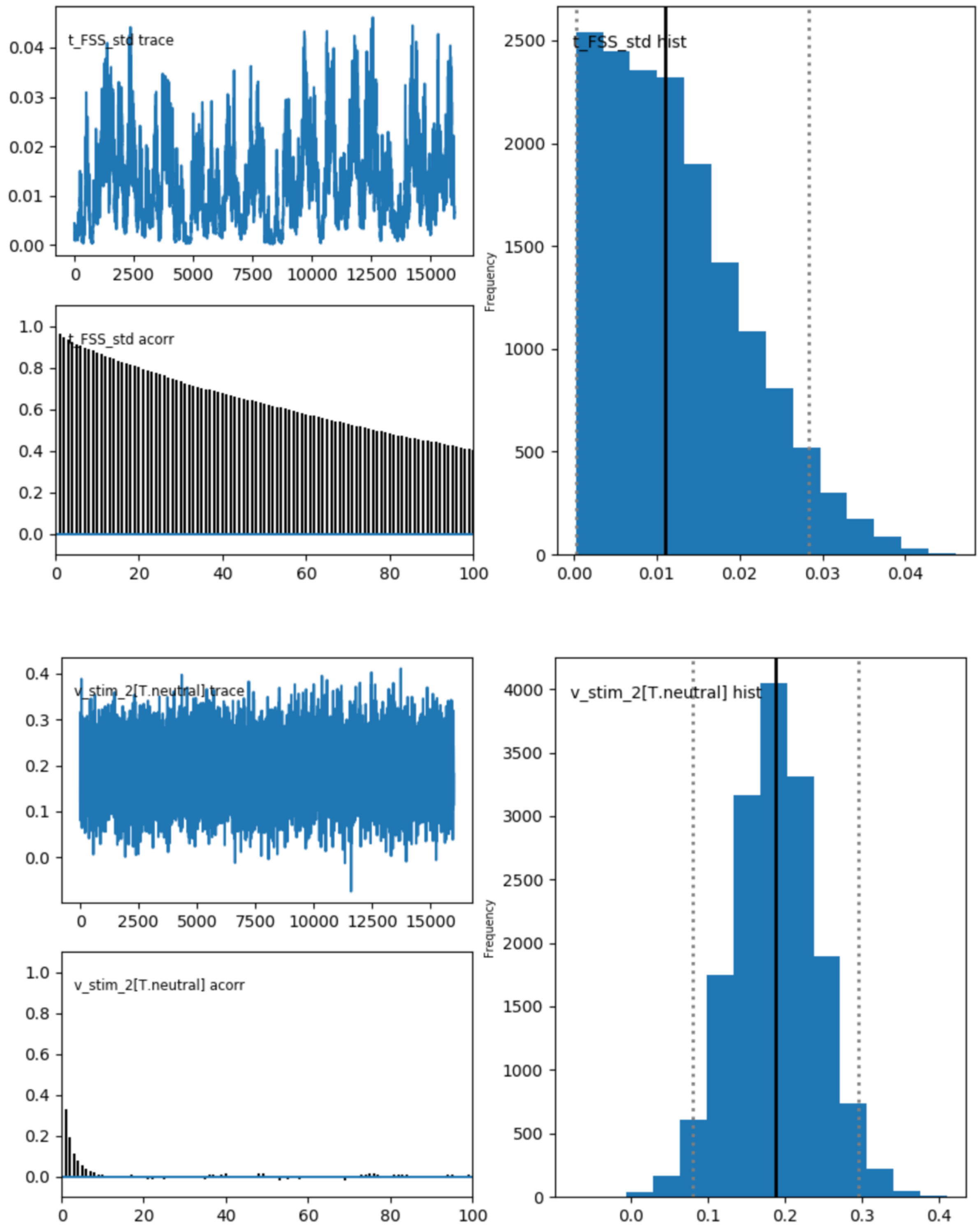


**Supplementary Fig. 4 (cont.).** Traces, autocorrelations and histograms of all parameters included in the hDDM regression model.

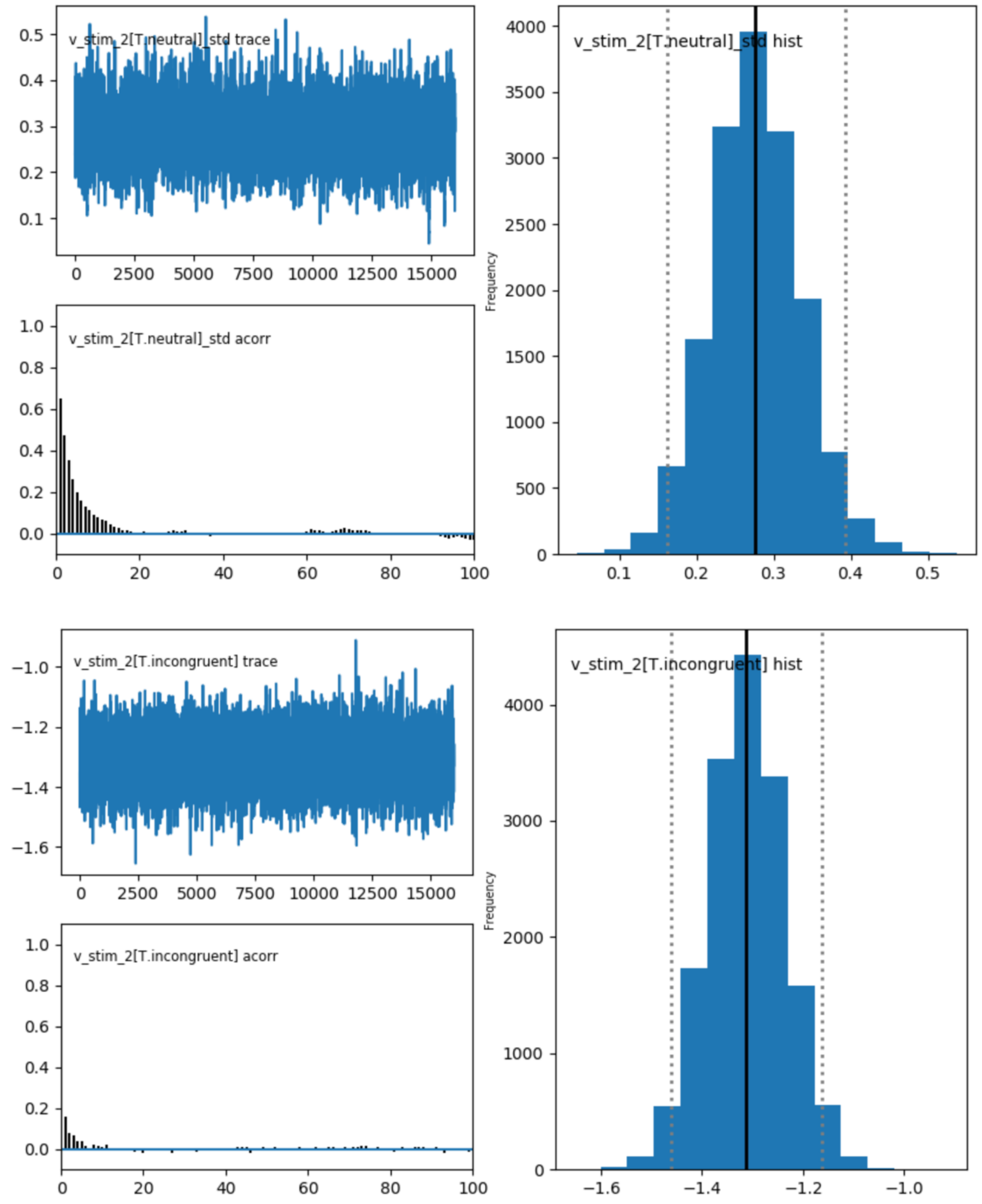


**Supplementary Fig. 4 (cont.).** Traces, autocorrelations and histograms of all parameters included in the hDDM regression model.

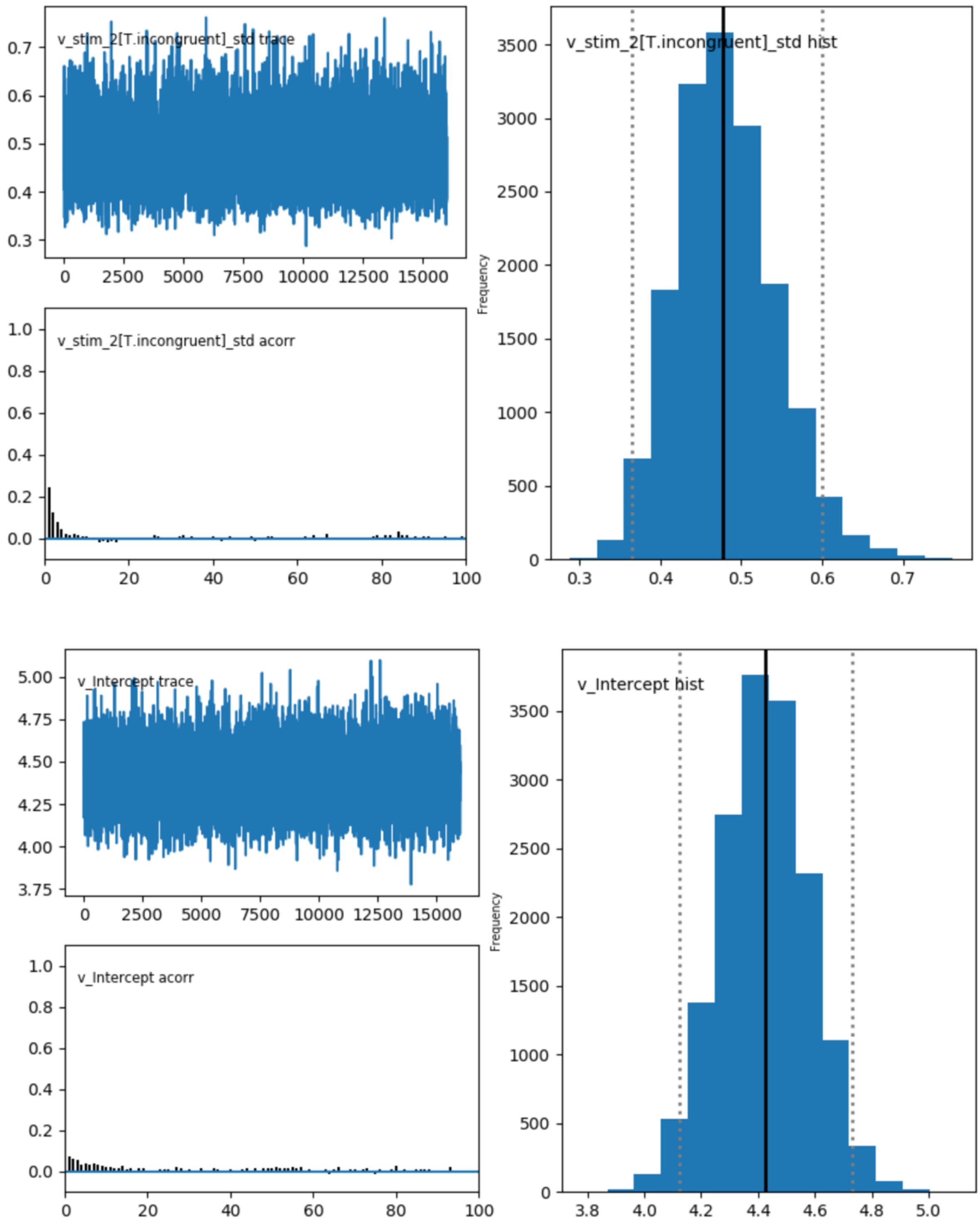


**Supplementary Fig. 4 (cont.).** Traces, autocorrelations and histograms of all parameters included in the hDDM regression model.

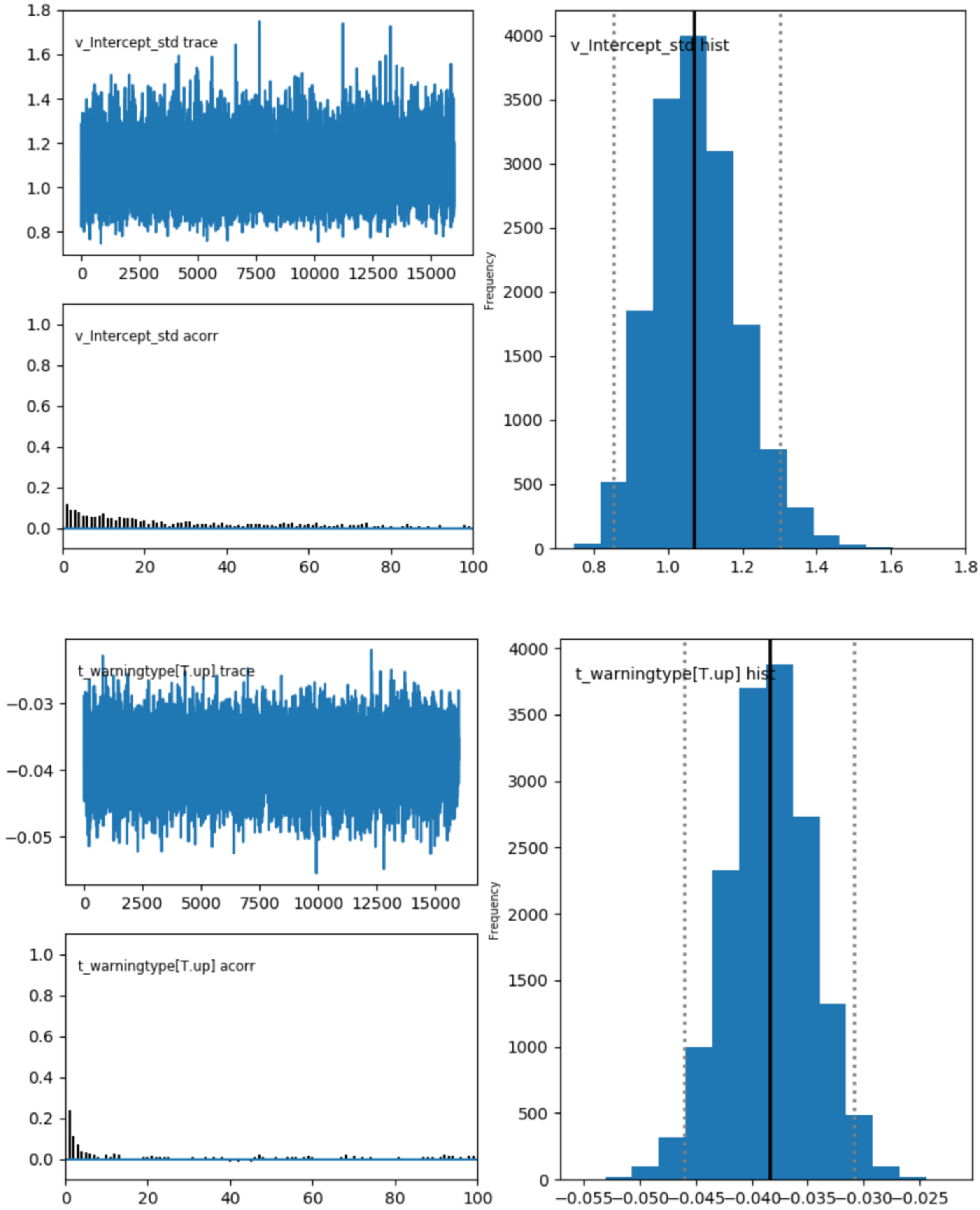


**Supplementary Fig. 4 (cont.).** Traces, autocorrelations and histograms of all parameters included in the hDDM regression model.

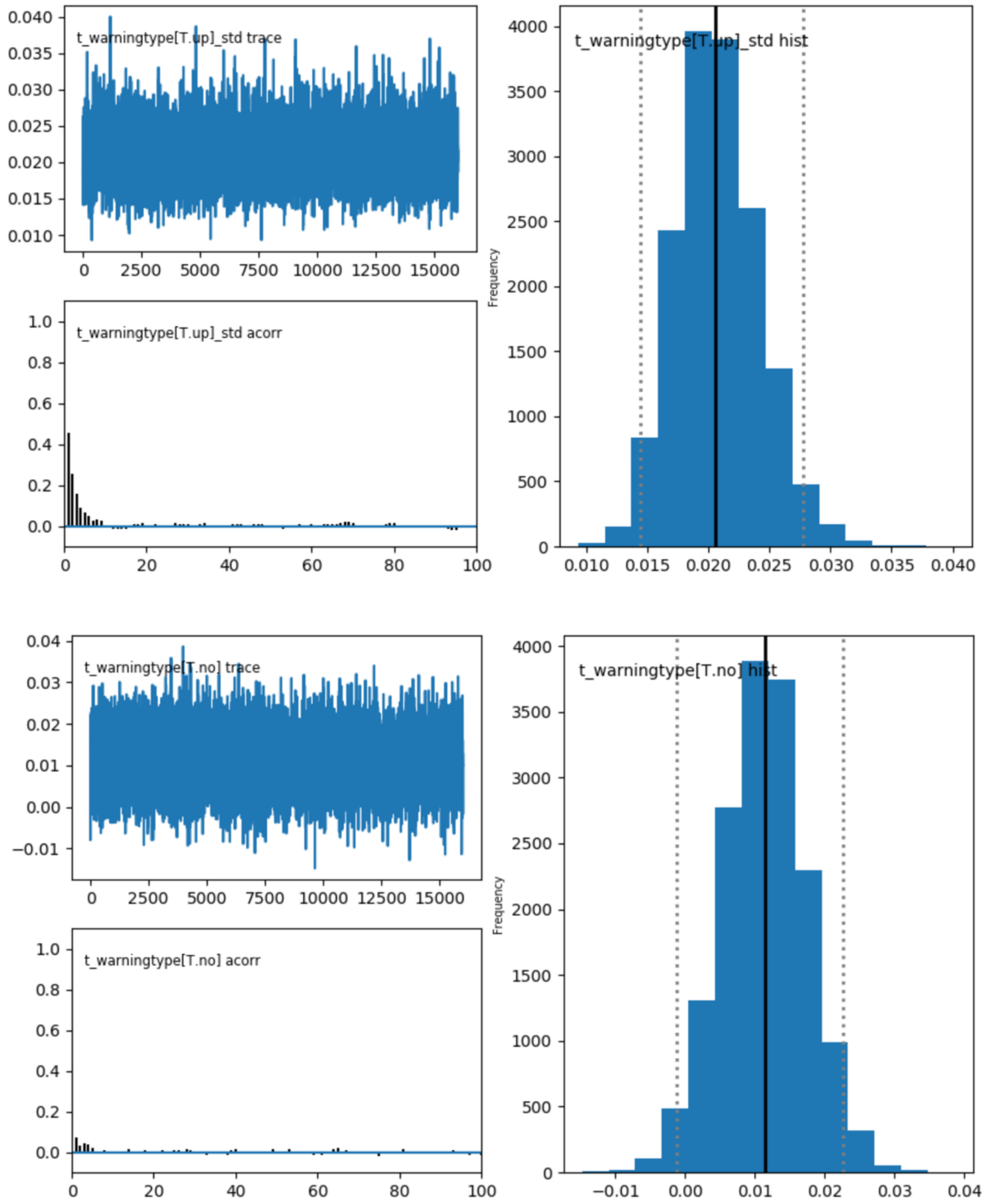


**Supplementary Fig. 4 (cont.).** Traces, autocorrelations and histograms of all parameters included in the hDDM regression model.

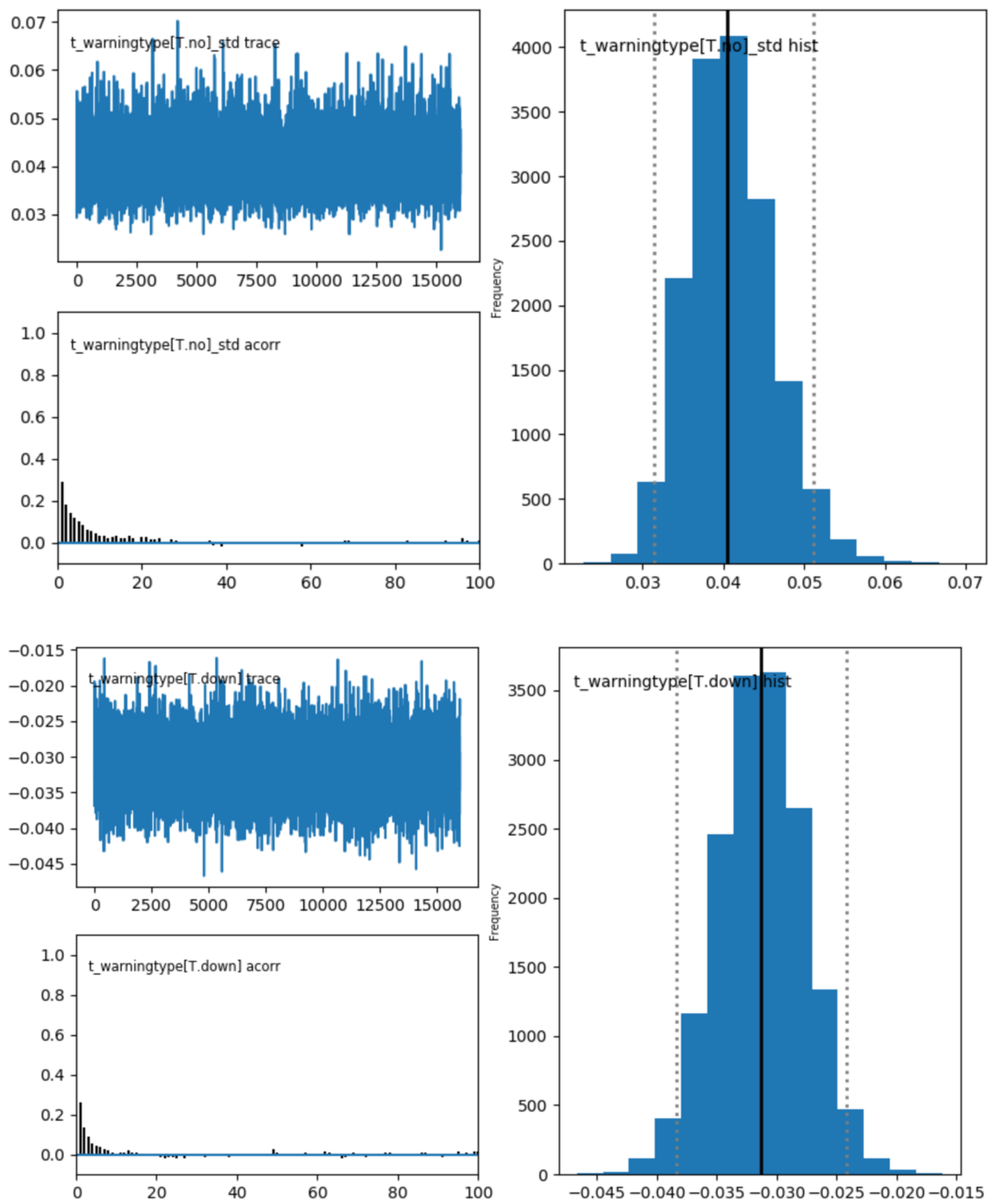


**Supplementary Fig. 4 (cont.).** Traces, autocorrelations and histograms of all parameters included in the hDDM regression model.

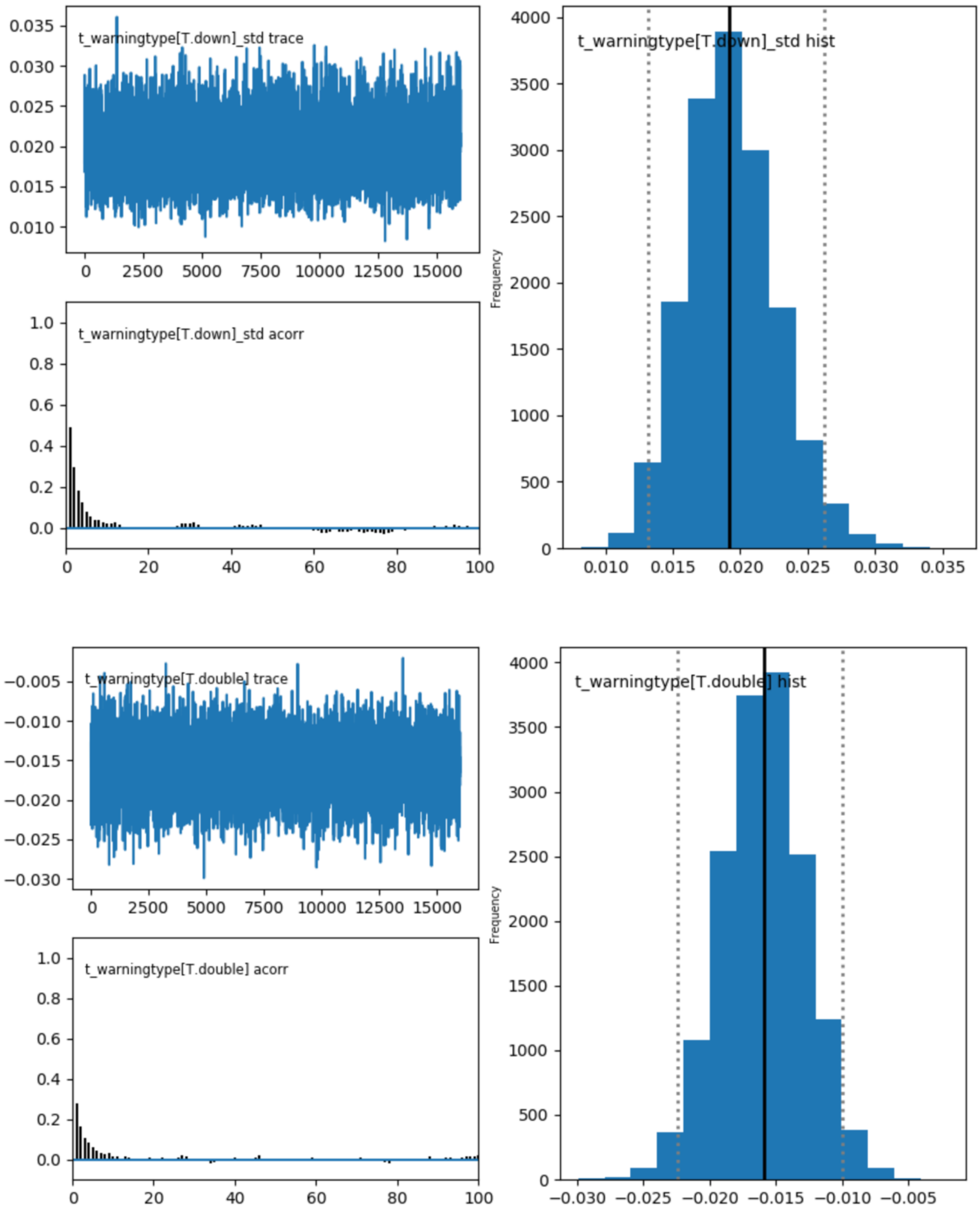


**Supplementary Fig. 4 (cont.).** Traces, autocorrelations and histograms of all parameters included in the hDDM regression model.

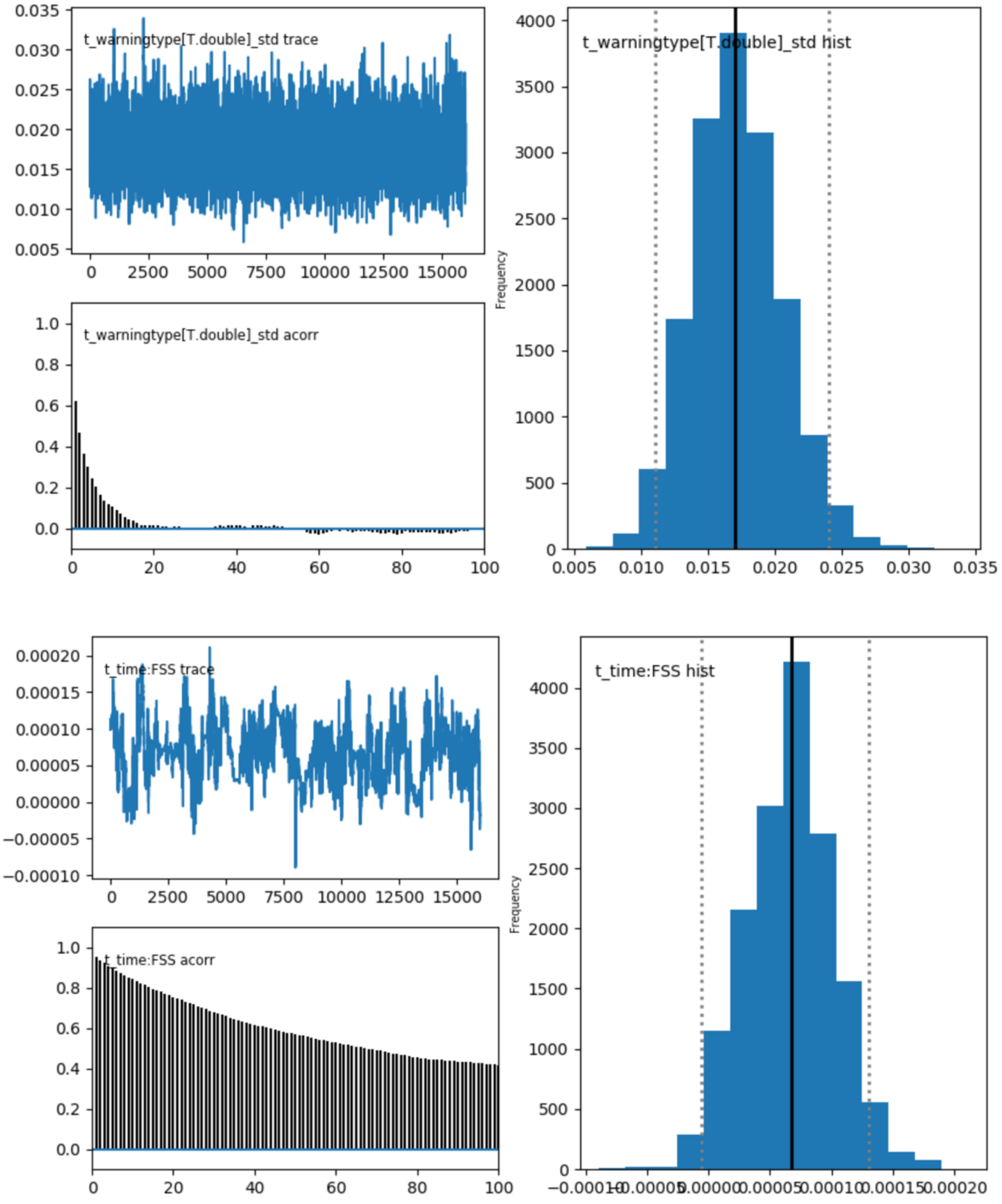


**Supplementary Fig. 4 (cont.).** Traces, autocorrelations and histograms of all parameters included in the hDDM regression model.

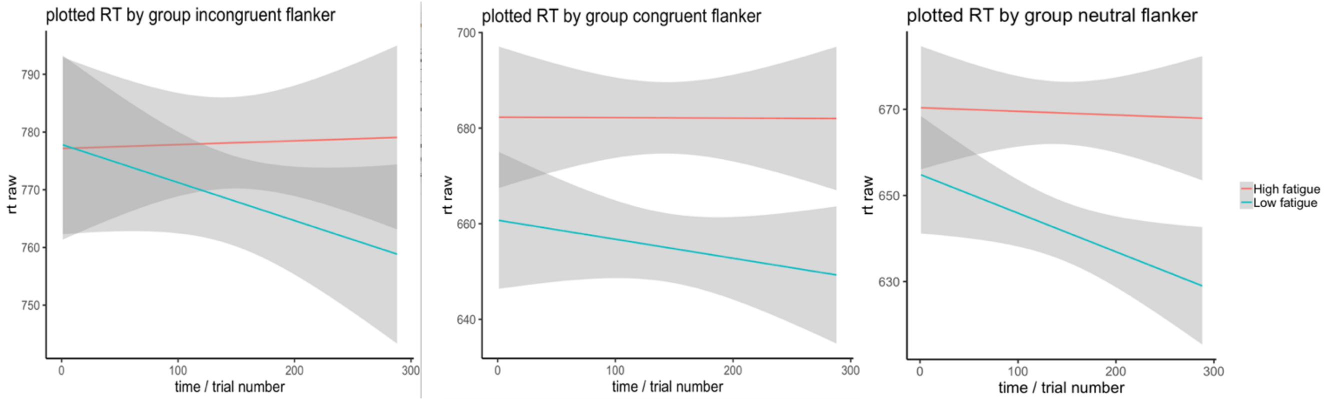


**Supplementary Fig. 5.** Raw RTs per target condition plotted as a function of trial number and group (high vs low fatigue, based on a median split).
